# Inhibition of TFIIH translocase reveals a novel lncRNA regulatory network at the BRCA1 locus

**DOI:** 10.1101/2025.03.11.642677

**Authors:** Samantha Cruz-Ruiz, Raphael Vidal, Mayra Furlan-Magaril, John T. Lis, Mario Zurita

**Affiliations:** Departamento de Genética del Desarrollo y Fisiología Molecular, Instituto de Biotecnología, Universidad Nacional Autónoma de México Av. Universidad 2001, Chamilpa, Cuernavaca Morelos, 62250. México; Department of Molecular Biology and Genetics, Cornell University Ithaca, NY 14853, USA; Departamento de Genética Molecular, Instituto de Fisiología Celular, Universidad Nacional Autónoma de México, Mexico City, México

**Keywords:** lncRNA/Promoters/Stress/Transcription

## Abstract

**Background:** Regulation of gene transcription is an essential early response to a broad variety of cellular stress. Paradoxically, pharmacological inhibition of an RNA polymerase II transcription initiation factor, XPB, induces a stress response that overactivates transcription of a subset of transcripts, including lncRNAs. However, the mechanisms enabling selective transcription under global initiation blockade, and the regulatory contribution of lncRNAs, remain unclear.

**Results:** Genome-wide assays in human cells treated with triptolide, which disrupts XPB translocase activity within TFIIH, revealed that a subset of genes continues to be transcribed independently of XPB activity, driven by thermodynamically unstable promoters. These promoters account for paradoxical transcriptional upregulation during global transcriptional inhibition. Among XPB-resistant transcripts, we identified previously uncharacterized regulatory long non-coding RNAs—*TILR-1*, *TILR-2*, and *LINC00910*—that are conserved in primates and display interdependent expression. Together, these lncRNAs form a coordinated regulatory module that modulates transcription at the *BRCA1* locus. Disruption of expression of these lncRNAs promotes cell proliferation and survival, phenocopying reduced expression of *BRCA1* locus genes. This lncRNA network is also activated by distinct transcriptional inhibitors indicating a shared transcriptional stress-responsive mechanism.

**Conclusions:** Unstable promoters sustain transcription in the absence of XPB, defining an XPB-independent transcriptional program that includes a transcriptional stress-responsive lncRNA regulatory network. This mechanism enables selective gene activation during global transcriptional inhibition, with the *BRCA1* locus representing one example, and suggests a broader transcriptional switch involved in cellular adaptation to transcriptional stress.

## Background

Transcription is a vital process for cells to regulate their growth and differentiation, as well as to respond rapidly to environmental changes and different types of stresses (1). RNA Polymerase II (RNA Pol II) transcription is regulated by the binding of transcription factors (TFs) to proximal and distal DNA elements, facilitating the recruitment and assembly of the pre-initiation complex (PIC) and then the productive elongation complex (2,3). Among the components of the PIC, the general transcription factor TFIIH is crucial for most transcription initiation, as its subunit XPB functions as a DNA translocase/helicase essential for DNA unwinding at the transcription start site (TSS), enabling RNA Pol II to initiate RNA synthesis, and hence, considered essential to transcription itself (4). A previous study showed that treating the oncogenic cell line MCF10A-ER-Src with the TFIIH inhibitors THZ1 and Triptolide (TPL) unexpectedly caused the upregulation of a group of genes, including long non-coding RNAs (lncRNAs) (5). However, the mechanisms underlying this paradoxical transcriptional activation and the functional role of these transcripts remain poorly understood.

LncRNAs, defined as transcripts longer than 500 nucleotides that do not have the potential to be translated into proteins, have been broadly studied in a variety of cellular contexts (6–9). Particularly, many examples of lncRNAs have been described as transcriptional regulators in development and diseases, such as cancer, playing roles at different levels of the transcriptional process (10–20). Moreover, regulatory lncRNAs have been studied under a variety of stresses such as hypoxia, energy stress, and DNA damage, among others (21–26). Nevertheless, how these lncRNA genes function during transcriptional stress caused by transcriptional inhibitors has not been broadly explored.

In this work, we describe the transcriptional stress response in the K562 cell line caused by TPL, using the genome-wide assays RNA-seq, PRO-seq, CUT&Tag of the activation histone mark H3K27ac, and ChIP-exo for the TFIIH subunit XPB. Using these strategies, we identify a group of TPL resistant genes that have thermodynamically unstable promoters, bypassing the XPB translocase activity. To gain a better understanding of this transcriptional stress response, we further characterized three long non-coding RNAs that are consistently upregulated in response to TPL treatment. The two novel lncRNAs, *<u>T</u>riptolide Induced <u>L</u>ong non-coding <u>R</u>NA* (*TILR*) –1 and –2, as well as the previously annotated *LINC00910*, form a coordinated regulatory module associated with the nearby locus containing the genes *BRCA1*, *NBR2*, *LOC101929767*, and *NBR1* (*BRCA1* locus). Finally, we explore the interdependence between these lncRNAs and the *BRCA1* locus, and how they bypass XPB inhibition to respond to this form of stress. Our findings define a subset of XPB-independent genes driven by unstable promoters and uncover a stress-responsive lncRNA regulatory network, providing genome-wide insights into selective transcription during global transcriptional inhibition.

## Results

### Triptolide global inhibition of transcription initiation leads to the accumulation of a subset of genomic transcripts

To investigate the molecular response caused by TPL in K562 cells, we first evaluated the cell survival, entry to apoptosis, and effects on the cell cycle following 24– and 48-hours treatments using flow cytometry. Our results show that this cell line is susceptible to cell death via apoptosis caused by transcriptional inhibition, with around 60% of cells entering apoptosis, and 50% of the cell population dead after 48 hours of 125 or 500 nM TPL treatments (Additional File 1: Fig. S1A-C). Additionally, the cell cycle was completely blocked at 24 hours of treatment (Additional File 1: Fig. S1D). These results are consistent with the effects of similar TPL treatments previously reported in other cell lines (5,27).

We then evaluated the effect of this drug after shorter treatments. When treating cells with TPL concentrations ranging from 125 nM to 1 μM for 4 and 8 hours, we found that cell survival was not affected, and cells did not enter apoptosis (Additional File 1: Fig. S2A-B). However, total RNA Pol II levels were reduced by at least 25% in the mildest treatment and up to around 55% in the rest of the conditions (Additional File 1: Fig. S2C). Based on these results, we determined that while prolonged TPL treatment is lethal, K562 cells remain viable under short-term exposure, allowing us to study the immediate transcriptional effects caused by inhibition of the TFIIH translocase activity.

To gain an insight into the response generated by this transcriptional inhibition, we performed RNA-seq experiments in K562 cells treated with 250 nM for 4 hours (Fig. 1A and Additional File 1: Fig. S2D). Differential expression analysis showed that most of the transcriptome was unaffected under these conditions (Additional File 1: Fig. S2E and Additional file 2). However, around 7.4% of transcripts were downregulated, while 8.8% were preferentially accumulated under TPL treatment (Additional File 1: Fig. S2E). This is consistent with a previous report using a similar TPL treatment (5). Among these transcripts, we detected 212 lncRNAs showing increased accumulation in response to this stress (Fig. 1B). Gene ontology (GO) analysis indicated that downregulated genes were enriched in development and differentiation functions, whereas upregulated transcripts were enriched in functions related to extracellular matrix organization, motility, and anion transport (Fig. S2F and Additional file 3). Given that lncRNAs are known to function as important regulators under different stress conditions (24–26), we further analyzed the upregulated lncRNAs using LncSEA 2.0, a comprehensive tool for functional enrichment analysis of lncRNAs integrating diverse datasets and annotations (28). Many of these lncRNAs were associated with survival, cancer phenotypes and hallmarks, among other categories (Fig. S2G and Additional file 3). These results indicate that a significant portion of transcripts that are preferentially accumulating after transcriptional inhibition by TPL correspond to lncRNAs potentially involved in important biological functions.

**Fig 1.**
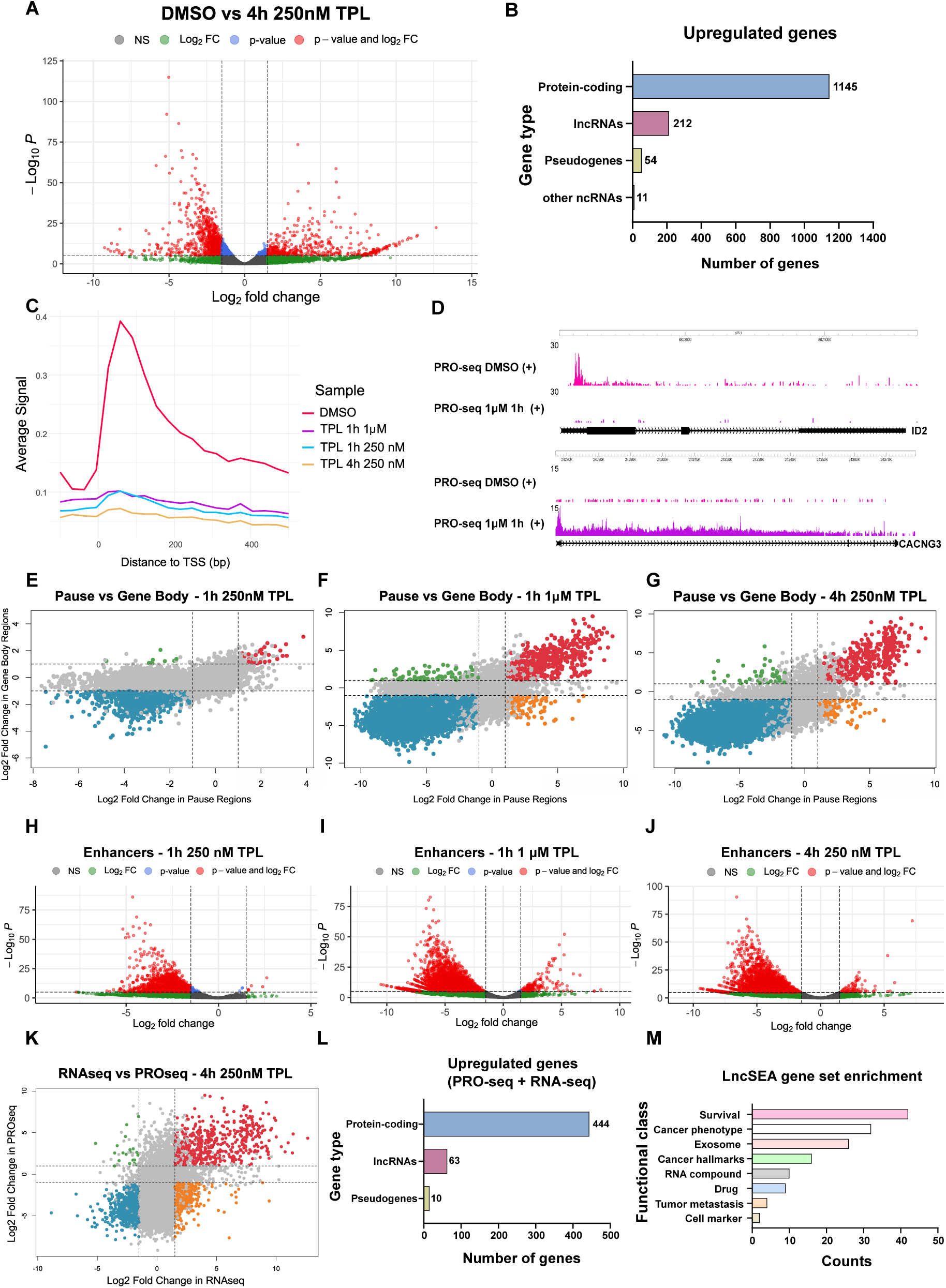
Triptolide treatment induces the overexpression of many protein-coding genes, lncRNAs, and cis-regulatory elements. **A** RNA-seq volcano plot of K562 cells treated with DMSO (Vehicle) or 250 nM TPL for 4 hours (n = 3 biological replicates per condition). Differentially expressed genes (Red dots) are those with a padj < 0.05 and fold change ≥ 1.5 (upregulated) or ≤ 1.5 (downregulated). **B** Gene type classification of the upregulated genes. **C** Metagene analysis of average PRO-seq signal at expressed genes around TSS upon TPL indicated treatments (n = 2 biological replicates per condition). Metaplot data is shown as an average signal in 10-nt bins. **D** Genome Browser image of examples of a downregulated gene (ID2, top) and an upregulated gene (CACNG3, bottom) upon 1h 1 μM of TPL treatment. **E-G** PRO-seq data showing the expression on pause region (x-axis) plotted against gene body region (y-axis) for TPL treatments of **E** 1h 250 nM, **F** 1h 1 μM, and **G** 4h 250 nM. Red dots = upregulated genes; blue dots = downregulated genes; orange dots = highly paused genes; green dots = high levels of expression in gene body only; grey dots: not significantly differentially expressed genes **H-J** PRO-seq volcano plots showing the differential expression of enhancers upon TPL treatments of **H** 1h 250 nM, **I** 1h 1 μM, and **J** 4h 250 nM. **K** Comparison between RNA-seq data (x-axis) and PRO-seq 4h 250 nM TPL treatment data (y-axis). Red dots = upregulated genes in both conditions; blue dots = downregulated genes in both conditions; orange dots = more accumulated genes only in RNA-seq data; green dots = genes significantly upregulated only in PRO-seq data. **L** Gene type classification of the upregulated genes found in both RNA-seq and PRO-seq 4h 250 nM data sets. **M** LncRNAs gene enrichment of the upregulated lncRNAs found in the RNA-seq and PRO-seq data using the software LncSEA 2.0.

Since RNA-seq cannot distinguish whether transcript accumulation is due to enhanced stabilization or transcriptional upregulation, we performed nascent RNA synthesis analysis using precision run-on sequencing (PRO-seq) under three different TPL treatments: 1h 250 nM (mildest treatment), 1h 1 μM (short but a high concentration treatment), and 4h 250 nM (longest treatment with a mild concentration) (Additional File 1: Fig. S3A-B). We found that transcription is highly affected in all the treatments (Fig. 1C). To be able to categorize the downregulated and upregulated transcripts, we plotted the fold changes in the pause regions versus the fold changes in the gene bodies, along with their respective p-values. Genes with significant increases in both pause region and gene body were considered upregulated, while genes with significant decreases in both pause region and gene body were considered downregulated. Examples of a highly downregulated and upregulated gene are presented in Fig. 1D. In the mildest treatment – 1h 250 nM, transcription is highly affected in most of the genes at the pause region, but it is not enough to cause a significant downregulation in the gene body (Fig. 1E, Additional File 1: Fig. S3C, and Additional file 4). This treatment also rapidly affected the expression of enhancers (Fig. 1H and Additional file 4), that were defined using the program dREG (29). Under this low TPL condition, only a few genes and enhancers actively responded to this transcriptional stress.

Analyzing the two remaining data sets corresponding to the 4h 250 nM and 1h 1 μM treatments, we observed a more pronounced transcriptional inhibition at both the promoter-proximal pause regions and gene bodies. Nevertheless, a significant number of genes (Fig. 1F-G, Additional File 1: Fig. S3D-E and Additional file 4) and enhancers (Fig. 1I-J and Additional file 4) were upregulated in response to these treatments. GO analysis revealed that downregulated genes were enriched in ribosome biogenesis, rRNA processing, and chromosome organization functions (Additional File 1: Fig. S3F-G and Additional file 5). For the upregulated genes, in the 1h 250 nM and 1h 1 μM treatments, we were not able to identify a functional category. However, for the 4h 250 nM dataset there was an enrichment in regulation of amine transport and receptor ligand activity, among other functions (Additional File 1: Fig. S3H and Additional file 5).

We further compared the results from 4h 250 nM TPL PRO-seq data to the corresponding RNA-seq and found that 517 genes are upregulated in both datasets, indicating that these genes that are more transcriptionally active give rise to stable RNAs (Fig. 1K). GO analysis of these shared genes showed similar results as the ones mentioned before (Additional File 1: Fig. S3I). Among the shared upregulated genes, 63 corresponded to lncRNAs (Fig. 1L), and enrichment analysis revealed that these lncRNAs are also related to survival, cancer phenotypes and hallmarks, among other categories (Fig. 1M and Additional file 5). By integrating these data sets, we determined a core of lncRNAs and protein encoding genes that are potentially implicated in the transcriptional stress response induced by TPL.

### XPB-independent promoters are less stably base-paired

To gain insight into the possible mechanism by which certain transcripts bypass the global transcriptional inhibition caused by TPL, we first examined changes in the histone mark H3K27ac. H3K27ac is a general activating mark for promoters and enhancers, that has been shown previously to require active RNA Pol II for its deposition (30). To determine whether TPL-induced upregulation of genes and enhancers triggers an increase in acetylation, we performed H3K27ac CUT&Tag experiments in K562 cells treated with 4h of 250 nM TPL (Additional File 1: Fig. S4A-B and Additional file 6). Analysis of the genomic distribution of differentially enriched regions showed that most downregulated acetylation upon transcription inhibition is localized in distal intergenic regions, introns, and promoters (Additional File 1: Fig. S4C and Fig. S4E-F). However, upregulated acetylation is in regions that were almost exclusively found at promoters and the first exon of the genes (Additional File 1: Fig. S4D and Fig. S4G). Comparison with PRO-seq and RNA-seq data from the same treatment revealed no correlation between gene expression and H3K27ac deposition under these conditions (Additional File 1: Fig. S4H-I). Intriguingly, all but one of the promoters showing increased acetylation correspond to downregulated genes. GO analysis indicated that this group is enriched in RNA metabolism functions, whereas genes showing decreased acetylation are associated with differentiation and proliferation pathways (Additional File 1: Fig. S4J and Additional file 6). These results indicate that changes in H3K27ac deposition do not explain the upregulation of genes under TPL treatment.

Because H3K27ac mark does not explain the resistance of some genes to TPL, we looked for other characteristics to gain insight into the mechanistic differences between the downregulated, upregulated, and unchanged genes upon TPL treatment. Past studies have shown that promoters with a DNA content that can be more easily melted do not require TFIIH translocase (XPB) activity (31,32). We hypothesized that the failure to inhibit certain promoter by TPL may be due to their less-stably base-pairing properties. Therefore, we performed GC-content analysis and found that the downregulated promoters have higher GC content percentage than the upregulated or unchanged promoters across all tested conditions (Fig. 2A-C). Moreover, by calculating the DNA duplex free energy using the nearest-neighbour method (33), we found that promoters of unchanged and upregulated genes are less-stably base-paired than downregulated promoters (Fig. 2D-F). These results indicate that promoters resistant to TPL-mediated transcriptional inhibition are less stably paired. Our findings align with previous studies showing that less stable promoters can be transcribed *in vitro* and in yeast independently of the TFIIH subunit XPB activity (31,32,34), the specific target of TPL.

**Fig 2.**
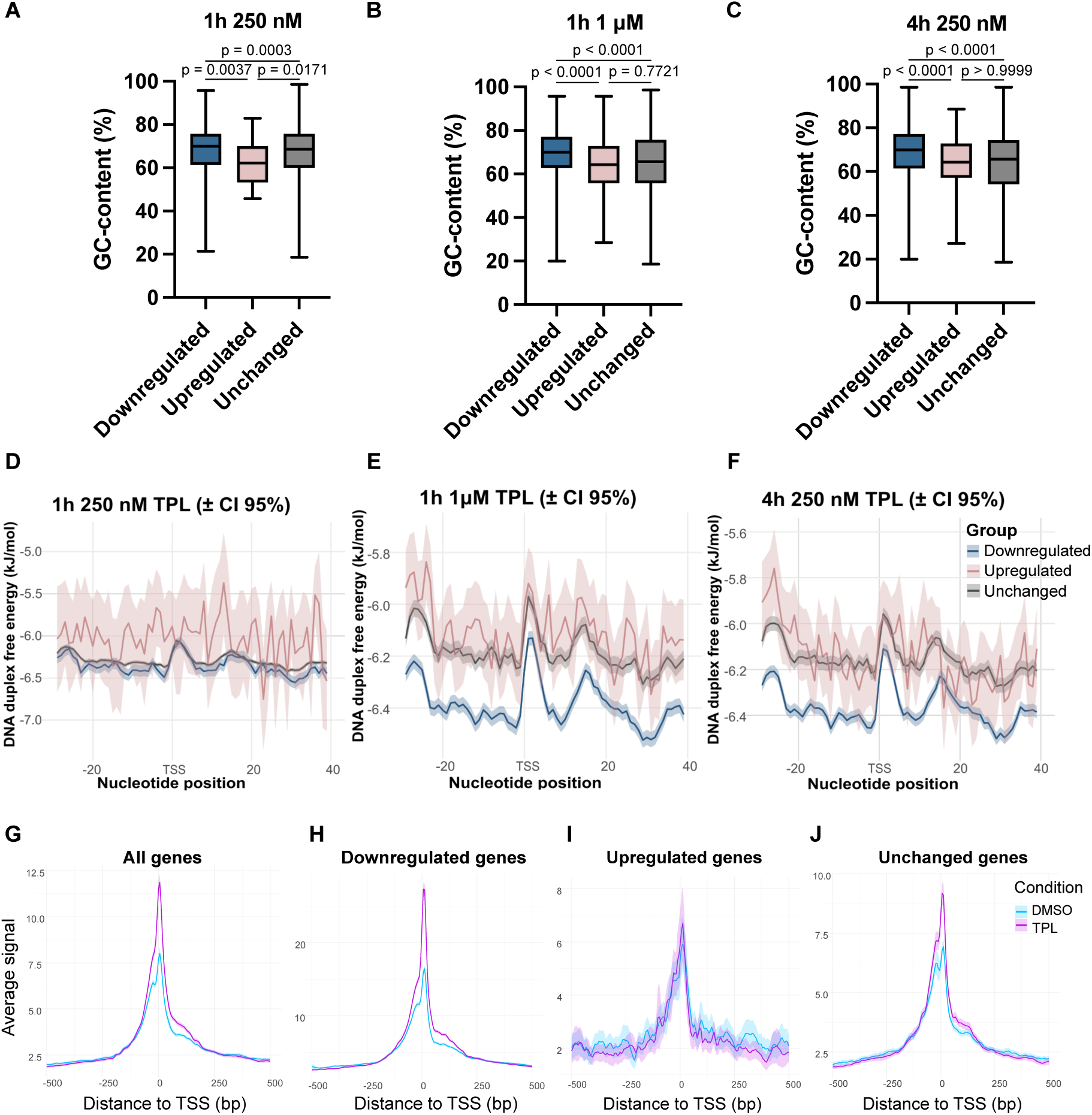
Promoter instability and XPB binding correlates with the TPL-induced downregulation and upregulation of a set of genes. **A-C** GC-content analysis from all the downregulated, upregulated, and unchanged promoters obtained from the **A** 1h 250 nM, **B** 1h 1 μM, and **C** 4h 250 nM PRO-seq analysis. Exact p-value is indicated above (Krustal-Wallis test). **D-F** DNA duplex free energy analysis performed from the same promoters used in the G-C content analysis. **G-J** Metagene analysis of XPB positioning in cells treated with DMSO or 1h 1 μM of TPL around the TSS of **G** all genes, as well as **H** downregulated, **I** upregulated, and **J** unchanged genes. Data is shown as an average signal in 10-nt bins.

To analyze if XPB occupancy changes upon TPL treatment, we performed ChIP-exo experiments in K562 cells treated with DMSO or 1h of 1 μM TPL. Meta-analysis revealed that the XPB peak around TSS is more pronounced in TPL-treated cells compared to DMSO controls (Fig. 2G). This pattern is consistent with previous observations in TPL-treated *Drosophila* cells (35) and suggests that XPB accumulates at promoters when its translocase activity is inhibited, thereby affecting transcription initiation by RNA Pol II. To explore this further, we examined XPB occupancy specifically at promoters of the downregulated, upregulated, and unchanged genes in response to TPL. Promoters of downregulated genes exhibited a marked increase in XPB binding (Fig. 2H). In contrast, upregulated promoters show no change in XPB occupancy between TPL– and DMSO-treated cells (Fig. 2I), while unchanged promoters display moderately elevated XPB levels (Fig. 2J). Therefore, these findings indicate that, in TPL-sensitive genes, inhibition of XPB ATPase activity leads to its accumulation at promoters, likely because its activity is required for dissociation from Pol II. However, in XPB-independent genes, transcription proceeds without changes in XPB binding, presumably because Pol II can enter elongation normally at these more easily melted promoters even in the absence of XPB activity.

### Two novel lncRNAs, *TILR-1* and *TILR-2*, along with *LINC00910*, are upregulated in response to global transcriptional inhibition by triptolide

To better understand the regulatory elements upregulated in response to TPL in PRO-seq datasets, we focused on two elements located near the lncRNA gene *LINC00910* that consistently responded to TPL treatment across all tested conditions (Fig. 3A, right; Additional file 4). Notably, *LINC00910* itself is also transcriptionally upregulated upon TPL treatment according to the PRO-seq data and shows preferential accumulation in the RNA-seq data (Fig. 3A, right). Both new elements were also classified as transcriptional regulatory elements (TREs) by the dREG software. Because these RNAs have not been previously described, we named them *Triptolide-Induced Long non-coding RNAs* (*TILR*)-1 and –2. These newly identified lncRNAs, together with *LINC00910*, are expressed across all examined human adult tissues (Additional File 1: Fig. S5)

**Fig 3.**
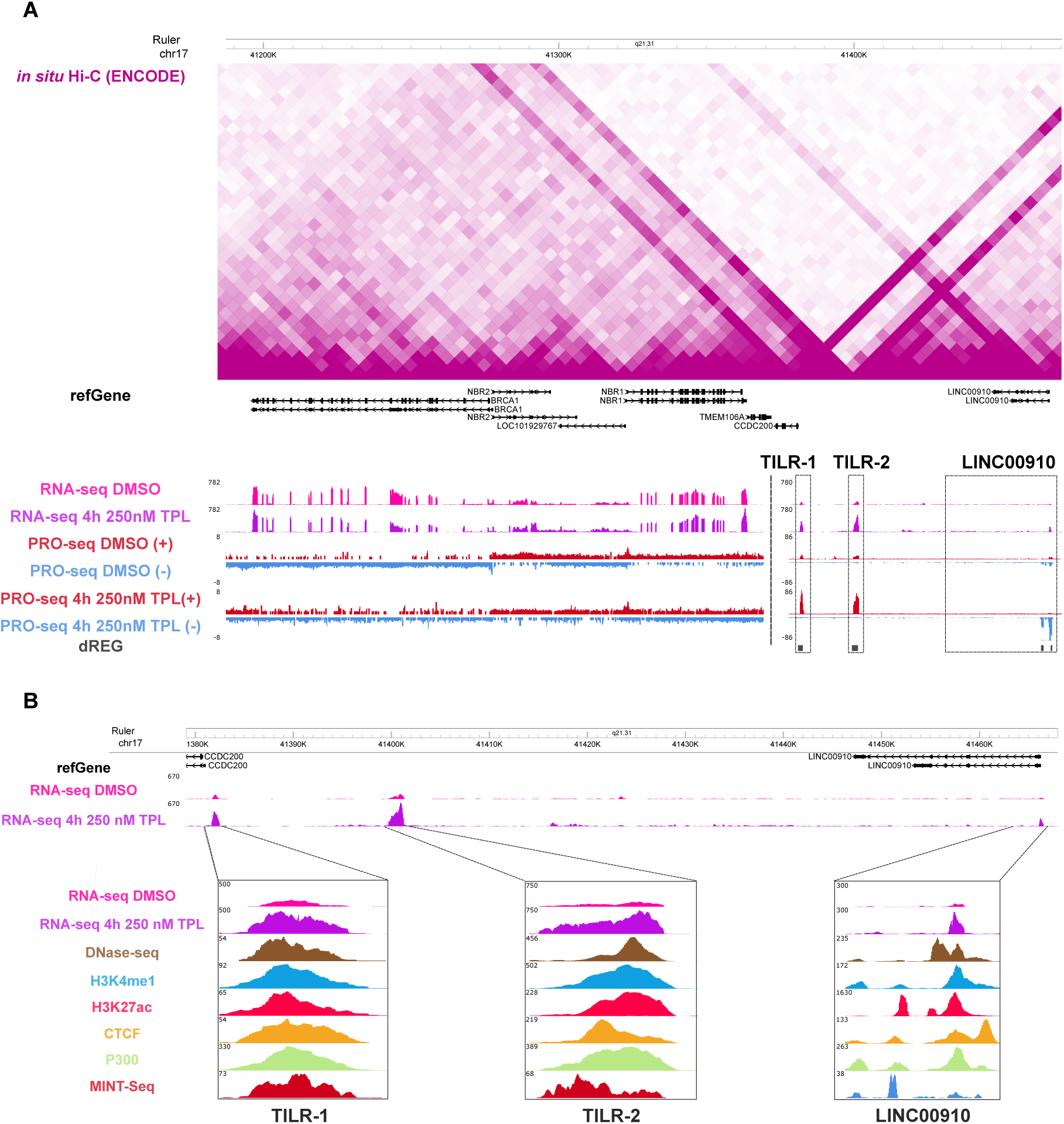
The discovery of the two novel lncRNAs *TILR-1* and *TILR-1* in chromosome 17q31. **A** Genome Browser image of the chromosome 17 spanning the *BRCA1* locus and the adjacent locus where the two novel lncRNAs *TILR-1*, *TILR-2*, and *LINC00910* reside. Hi-C data was retrieved from ENCODE. **B** Genome browser image of the region corresponding to *TILR-1*, *TILR-2*, and *LINC00910*. Shown are RNA-seq data in K562 cells treated with DMSO or 4h 250 nM TPL, and public data from ENCODE showing the chromatin accessibility in these elements (DNAse-seq), as well as the presence of the enhancer marks H3K27ac and H3K4me1, and the transcription factors CTCF and P300. MINT-seq data corresponding to the RNA modification m6A was downloaded from (37).

Since these elements were detected as potential regulatory elements, we explored *in situ* Hi-C data from K562 in the ENCODE portal (36) and observed that these three regions are physically connected to each other, as well as to the adjacent *BRCA1* locus, a tumor suppressor locus containing the genes *BRCA1*, *NBR2*, *NBR1*, and *LOC101929767* (Fig. 3A, top). These findings led us to further investigate the potential function of these elements during transcriptional stress induced by TPL treatment.

### Long non-coding RNAs *TILR-1*, *TILR-2*, and *LINC00910* have characteristics of regulatory elements

To further characterize the nature of these non-coding elements, we searched for genomic data available in ENCODE (36), UCSC Genome Browser, and Gene Omnibus Database, and analyzed whether regions display regulatory element signatures, particularly since *TILR-1* and *TILR-2* were identified as TREs in the PRO-seq analysis. Data from the UCSC Genome Browser show that these elements are not annotated in the Ensembl, are not ribosome-protected according to Ribo-seq data, and are conserved only in primates, among other features (Additional File 1: Fig. S6). Moreover, as shown in Fig. 3B and Additional File 1: Fig. S7A, these elements display hallmarks of active regulatory elements in K562 cells, including active transcription (38–40), DNase hypersensitive sites, high levels of H3K4me1 and H3K27ac, binding of multiple transcription factors (TFs) such as P300 and CTCF (41), and the presence of the m6A modification in the RNA, which is characteristic of long and stable enhancer RNAs (37). Importantly, these signatures are not limited to K562 cells but are also observed in MCF7 and MCF10A-ER-Src cell lines (Additional file 7).

Following these results, we evaluated the length of the *TILR-1* and *TILR-2* transcripts and found that *TILR-1* is at least 941 nt long (Additional File 1: Fig. S7B) and *TILR-2* is at least 1.3 kb long (Additional File 1: Fig. S7C). These RNAs also contain poly-A tails (Additional File 1: Fig. S7D) and lack an open reading frame along their sequences. These features are characteristic of a class of enhancers that produce lncRNAs rather than the typical bidirectional eRNA transcripts, and that are required for proper transcription of their target genes (37,41–44). The complete *TILR-1* and *TILR-2*, as well as their genomic coordinates are shown in Additional File 1: Fig. S8.

Next, we performed cellular fractionation experiments followed by RT-qPCR to determine the subcellular localization of *TILR-1*, *TILR-2*, and *LINC00910*. As shown in Fig. 4A-C and Additional File 1: Fig. S7E-G, these lncRNAs are highly enriched in the chromatin fraction in both control (DMSO) and TPL-treated conditions. Importantly, as controls for fractionation, RT–qPCR was performed using 7SL RNA as a cytoplasm-associated RNA positive control and the 45S rRNA precursor as a chromatin-associated RNA positive control (Additional File 1: Fig. S7H-I). Western blot analysis was also performed using GAPDH, Lamin B, and Histone H3 as cytoplasmic, nucleoplasmic, and chromatin-associated markers, respectively. (Additional File 1: Fig. S7J). To confirm this observation, we performed RNA fluorescence *in situ* hybridization (FISH) using probes against *TILR-2* and *LINC00910*. Probe specificity was validated using *TILR-2* and *LINC00910*-depleted cells (Additional File 1: Fig. S9A). We found that these lncRNAs accumulate in foci (Fig. 2D), and notably, the area of these foci increases upon TPL treatment (Fig. 4E-F and Additional File 1: Fig. S9B-C), suggesting that upregulated *TILR-2* and *LINC00910* RNAs are retained and accumulate within the same chromatin region. Importantly, we consistently observed between two and three foci per cell in most analyzed cells, as expected for this near-triploid cell line (Additional File 1: Fig. S9D-E). Treatment of K562 cells with different TPL concentrations and time points results in the upregulation of these elements under almost all the conditions (Additional File 1: Fig. S9F-H).

**Fig 4.**
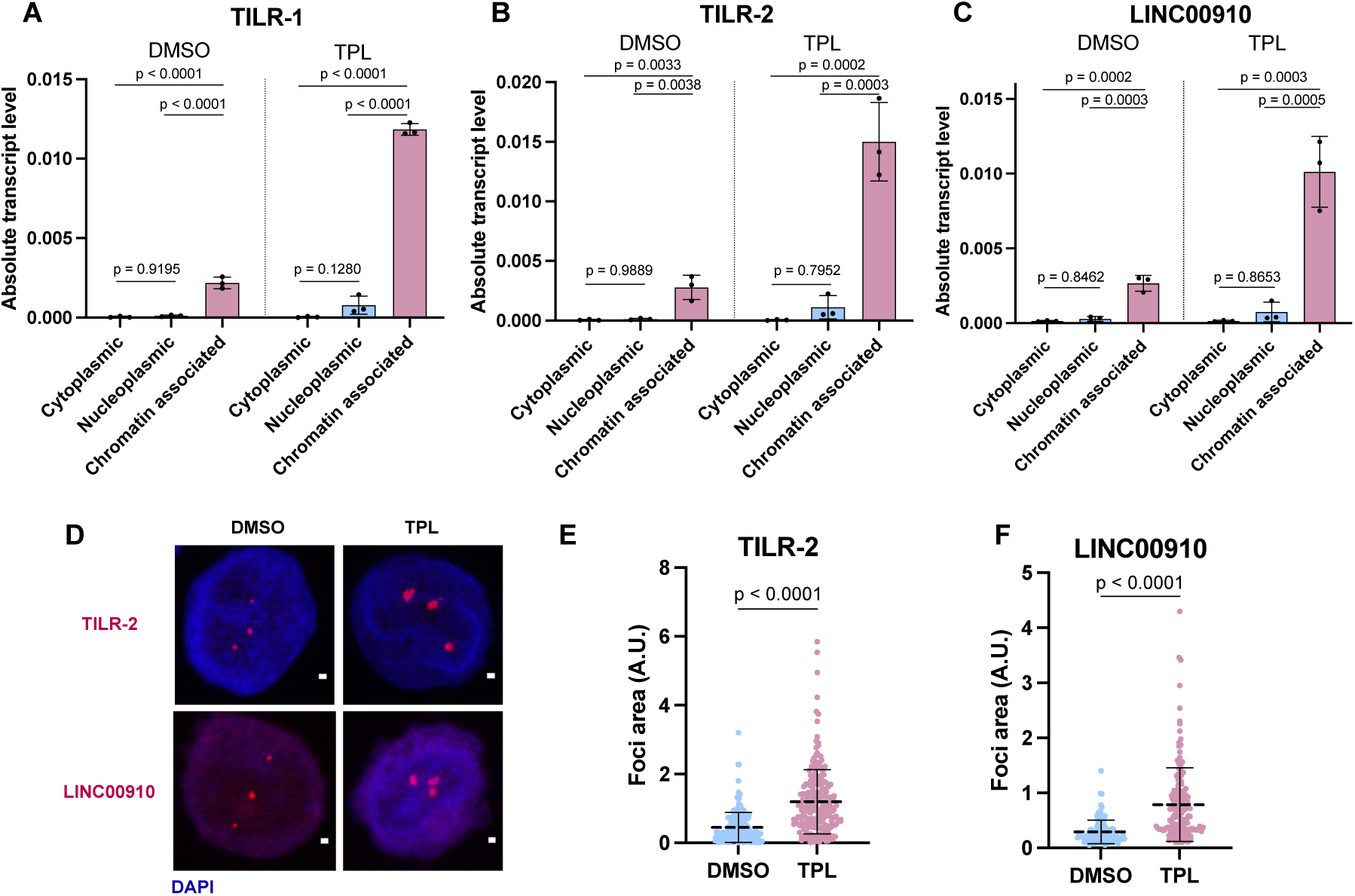
*TILR-1*, *TILR-2*, and *LINC00910*, present hallmarks of regulatory elements. **A-C** Cellular fractionation experiments followed by RT-qPCRs (absolute quantification) evaluating the expression of *TILR-1* **A**, *TILR-2* **B**, and *LINC00910* **C** in K562 cells treated with DMSO or 4h 250 nM TPL. Data represent mean ± SD of n = 3 biological replicates. Exact p-value is indicated above (one-way ANOVA test). **D** Representative images of *TILR-2* and *LINC00910* FISH in K562 cells treated with DMSO or 4h 250 nM TPL. Three biological replicates were performed and at least 80 cells per condition were analyzed. Scale bar = 1 μm. **E-F** Dot plot showing the quantification of **E** *TILR-2* and **F** *LINC00910* FISH foci area (A.U. = arbitrary units). Each dot represents the quantification of individual cells. Exact p-value is indicated above (Mann-Whitney test).

Finally, we tested their expression upon TPL treatment in MCF7, MDA-MB-231, and MCF10A-ER-Src cell lines, and observed that these lncRNAs are also upregulated in these cell types (Additional File 1: Fig. S9I-N). These results indicate that *TILR-1, TILR-2*, and *LINC00910* are potential regulatory elements producing long, stable RNA products across different cellular contexts in response to transcriptional stress.

### Expression of the lncRNAs *TILR-1*, *TILR-2*, and *LINC00910* is interdependent, and these elements regulate the expression of *BRCA1*, *NBR1*, *NBR2*, and *LOC101929767*

To investigate the potential function of these lncRNAs, we performed a variety of knockdown strategies. First, we conducted CRISPR interference (CRISPRi) experiments by generating K562 cell lines constitutively expressing dCas9 fused to the repressor domain KRAB (dCas9-KRAB) constitutively (Additional File 1: Fig. S10A). We then designed sgRNAs against *TILR-1*, *TILR-2*, and *LINC00910* to transfect them transiently. Forty-eight hours post-nucleofection, we measured the expression of each RNA and observed an interdependence effect among these three RNAs – the impairment of one influences the expression of the other two – under normal conditions, as well as upon 250 nM TPL treatment for 4 h, as shown in Fig. S10B-D.

Given that the KRAB domain can induce heterochromatinization near its target region, we generated another cell line expressing dCas9 alone to cause a transcriptional interference effect rather than a silencing effect (Additional File 1: Fig. S10A). Although the interference effect was less pronounced than silencing, we observed the same interdependence effect in expression of the three RNAs under normal conditions (Additional File 1: Fig. S10H-J). Interestingly, upon TPL treatment, we observed higher levels of variability in the expression of these RNAs (Additional File 1: Fig. S10H-J), suggesting that dCas9 interference can sometimes be overcome by RNA Pol II recruitment to upregulate these elements. Importantly, this effect is specific, as a nearby gene that is downregulated upon TPL treatment showed no altered response when we transfected the sgRNAs (Additional File 1: Fig. S10E and S10K). These results indicate that expression of *TILR-1*, *TILR-2*, and *LINC00910* is mutually dependent. Cell viability assays after 48 hours of incubation with DMSO or TPL showed no significant difference between the silenced cells and controls (Additional File 1: Fig. S10F-G and S10L-M).

Since these CRISPRi experiments do not allow us to distinguish whether transcription or the RNA product itself mediates this regulation, we performed knockdown using antisense oligonucleotides (ASOs), a widely used strategy for functional characterization of lncRNAs (10,45). We designed one ASO against each RNA (Additional File 1: Fig. S11A) and transfected them individually or pooled together. To avoid any effect of the ASOs on lncRNA transcription, the target sequences were designed far from their TSS (Additional File 1: Fig. S11A). An ASO against GFP was used as negative control. Six hours post-nucleofection, followed by 250 nM TPL treatment for 4h (Fig. 5A-C) or 1 μM TPL treatment for 1h (Additional File 1: Fig. S11B-D), allowed us to have better silencing effects. In both treatments, we observed results consistent with the previously described CRISPRi experiments, suggesting that the RNAs themselves mediate the interdependence of their expression.

**Fig 5.**
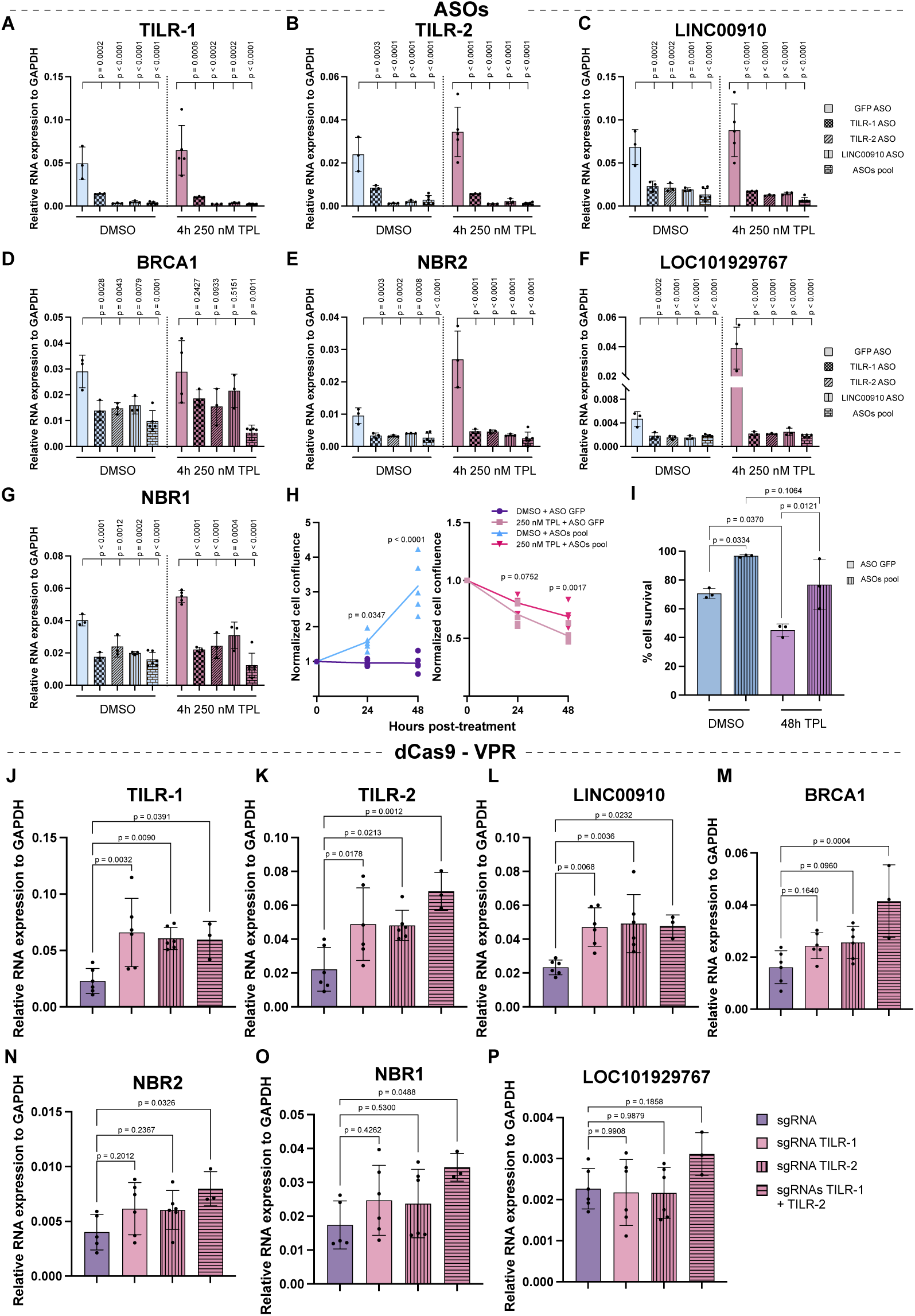
Expression of the lncRNAs *TILR-1*, *TILR-2*, and *LINC00910* is interdependent, and these elements regulate the expression of *BRCA1*, *NBR1*, *NBR2*, and *LOC101929767*. **A-G** RT-qPCR to measure the expression of **A** *TILR-1*, **B** *TILR-2*, **C** *LINC00910*, **D** *BRCA1*, **E** *NBR2*, **F** *LOC101929767*, and **G** *NBR1* 6 hours after nucleofection of ASOs for the knockdown of the lncRNAs *TILR-1*, *TILR-2*, and *LINC00910* + 4 hours of treatment with DMSO or 4h 250 nM TPL. As negative control, an ASO against GFP was used. Data represent mean ± SD of at least n = 3 biological replicates. Exact p-value is indicated above (one-way ANOVA test). **H** Cell proliferation of transfected cells with a pool of ASOs against TILR-1, TILR-2, and LINC00910, or against GFP as a control. These cells were treated with DMSO (left) or 250 nM TPL (right) for 24 and 48 hours, and cells were counted after these treatments. Data represent mean ± SD of n = 5 biological replicates. Exact p-value is indicated above (two-way ANOVA test). (**I**) Cell survival of transfected cells with a pool of ASOs against TILR-1, TILR-2, and LINC00910, or against GFP as a control. These cells were treated with DMSO or 250 nM TPL for 48 hours. Data represent mean ± SD of n = 3 biological replicates. Exact p-value is indicated above (one-way ANOVA test). **J-P** RT-qPCR to measure the expression of **J** *TILR-1*, **K** *TILR-2*, **L** *LINC00910*, **M** *BRCA1*, **N** *NBR2*, **O**, *NBR1* and **P** *LOC101929767*6 48 hours after nucleofection of dCas9-VPR + sgRNAs targeting TILR-1 or TILR-2 for overexpression. As negative control, sgRNA plasmid without gRNAs was used. Data represent mean ± SD of at least n = 3 biological replicates. Exact p-value is indicated above (one-way ANOVA test).

Next, we evaluated the effect on the potential target genes under these knockdown conditions. Expression of *BRCA1, NBR1, NBR2*, and *LOC101929767* genes was significantly reduced upon depletion of *TILR-1, TILR-2*, and *LINC00910* RNAs (Fig. 5D-G and Additional File 1: Fig. S11E-H), confirming that these lncRNAs act as positive regulators of these genes. Cellular proliferation assays revealed that knockdown of these lncRNAs increases cell proliferation after 48 hours (Fig. 5H), and cells are more viable under both DMSO and TPL conditions (Fig. 5I and Additional File 1: Fig. S11I) suggesting that these RNAs are down-modulators of proliferation and survival via regulation of their target genes.

To assess whether the reciprocal effect occurs under overexpression, we used the CRISPRa system with dCas9 fused to the transcriptional activator VPR (dCas9-VPR) to upregulate *TILR-1* and *TILR-2* in *cis*. We designed sgRNAs targeting *TILR-1* and *TILR-2*, transfected them individually or together, along with the plasmid containing the dCas9-VPR, and evaluated transcript levels 48 hours post-nucleofection. As shown in Figure 5J-L, we observed the same interdependence effect: overexpression of either *TILR-1* or *TILR-2* upregulates all three lncRNAs, including *LINC00910*. We then evaluated the effect on the *BRCA1* locus and found that *BRCA1*, *NBR1*, and *NBR2* are significantly upregulated when both sgRNAs targeting *TILR-1* and *TILR-2* are transfected simultaneously (Figure 5M-O). In the case of *LOC101929767*, a tendency toward upregulation was observed, but it did not reach statistical significance (Figure 5P). Although overexpression was mild, these data support the interdependence effect and the role of these lncRNAs in regulating the *BRCA1* locus.

ENCODE data show that the genomic regions of *TILR-1*, *TILR-2*, and *LINC00910* bind multiple transcription factors in the K562 cell line (Fig. 2B and Additional File 1: Fig. S7A). We therefore extended our analysis to examine TF binding near or directly at the bidirectional promoters of the four-gene cluster. Interestingly, several TFs that bind to *TILR-1, TILR-2*, and *LINC00910* also occupy sequences near the two bidirectional promoters—one driving the transcription of *BRCA1* and *NBR2*, and the other for *LOC101929767* and *NBR1* (Additional File 1: Fig. S11J). These results suggest that TFs binding to *TILR-1*, *TILR-2*, and *LINC00910* may also bind the bidirectional promoters within the *BRCA1* locus, potentially contributing to the coordinated regulation of this gene cluster.

### *TILR-1*, *TILR-2*, and *LINC00910* exhibit responses to a range of transcriptional inhibitors

Since these lncRNAs are overexpressed in different cell lines as a response to various concentrations of TPL, we aimed to test whether their upregulation is specifically due to inhibition of XPB or whether it also occurs when other factors involved in early transcription are inhibited. To address this, we analyzed public RNA-seq data from different cell lines treated with a variety of transcriptional inhibitors targeting distinct components of the transcription machinery, including XPB (TPL), CDK7 (THZ1), CDK9 (NVP-2), BRD4 (JQ1), and histone acetyltransferase activity (C646), all of which impact early transcriptional regulation transcription (46–50). We found that in many cases *TILR-1, TILR-2*, and *LINC00910* were upregulated in response to these treatments in different cell lines (Fig. 6A), We further analyzed public data from K562 cells treated with JQ1 and C646 and observed that these lncRNAs were also upregulated under these conditions (Fig. 6A). Finally, to determine whether this upregulation occurs in systems more comparable to physiological conditions, we analyzed public RNA-seq data from primary T-cells treated with Actinomycin D (51), and from two pancreatic cancer organoids (U049M15 and U049MAI) treated with the pro-drug CK21, a TPL analogue (52). In T-cells treated with Actinomycin D, only *TILR-1* was upregulated, whereas *TILR-2* and *LINC00910* showed no changes (Fig. 6A). In contrast, in the pancreatic cancer organoids treated with CK21 for 3, 6, 9, and 12 hours, all three lncRNAs were clearly upregulated after 9– and 12– hours of treatment (Fig. 6B-D). These results suggest that *TILR-1*, *TILR-2*, and *LINC00910* respond to transcriptional stress across multiple treatments, in various cell lines, and in 3D organoid cancer models, supporting their potential biological relevance in disease contexts. Moreover, these findings further support the model in which these lncRNAs may continue to be expressed due, at least in part, to the thermodynamically instability of their transcribed regions.

**Fig 6.**
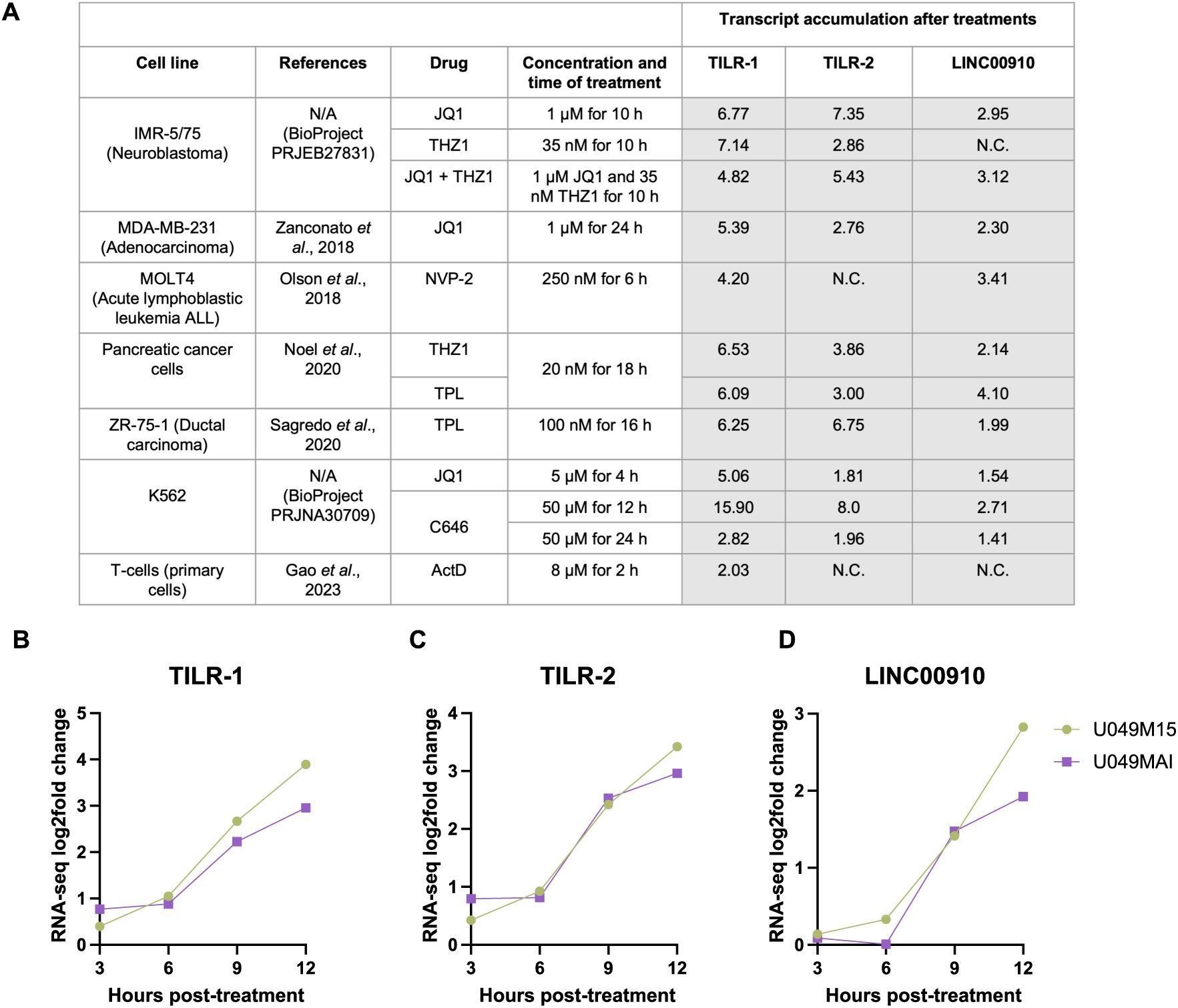
LncRNAs *TILR-1*, *TILR-2*, and *LINC00910* are upregulated upon treatment with other transcriptional inhibitors and under heat shock and arsenic-induced stress. **A** Summary of the expression of the lncRNAs *TILR-1*, *TILR-2*, and *LINC00910* in different cell lines treated with a variety of transcriptional inhibitors, according to public RNA-seq data. N.C = no change. **B-D** RNA-seq log2fold change values showing the expression of **B** *TILR-1*, **C** *TILR-2*, and **D** *LINC00910* in pancreatic cancer organoid originated from two patients (U049M15 and U049MAI). RNA-seq data was downloaded from the GEO number GSE225011.

### LncRNAs *TILR-1*, *TILR-2*, and *LINC00910* are interconnected with the *BRCA1* locus via DNA-DNA and RNA-DNA

Analysis of XPB occupancy along the *BRCA1* locus and the lncRNAs *TILR-1*, *TILR-2*, and *LINC00910* revealed that both bidirectional promoters within the *BRCA1* locus exhibit increased XPB binding (Additional File 1: Fig. S12A). In contrast, the genomic regions corresponding to *TILR-1* and *TILR-2*, as well as the promoter of *LINC00910*, behave as TPL-resistant loci, showing no significant change in XPB occupancy (Additional File 1: Fig. S12B). We hypothesize that *TILR-1*, *TILR-2*, and *LINC00910* counteract the sensitivity of *BRCA1* locus promoters to TPL, enabling an effective transcriptional response under stress conditions.

From public *in situ* Hi-C data, these elements are observed to physically interact with the *BRCA1* locus (Fig. 2A). To investigate whether the lncRNAs interact with the *BRCA1* locus genomic region, we analyzed public ChAR-seq (53) dataset from K562. We also generated a virtual 4C map from previously visualized *in situ* Hi-C data (54). The virtual 4C analysis (Additional File 1: Fig. S12C, top) highlights interactions between the lncRNAs and the *BRCA1* locus. ChAR-seq data (Additional File 1: Fig. S13C, bottom) indicate that *LINC00910* primarily interacts with the *BRCA1* locus DNA region. Based on these results, we propose that these three lncRNAs may form part of a transcriptional hub necessary for coordinated expression of this gene cluster containing the *BRCA1* locus. Although the precise mechanism of these RNA-DNA interactions remains unclear, we hypothesize that repetitive elements, particularly Alu sequences, may mediate these interactions (see Discussion).

### LncRNAs *TILR-1*, *TILR-2*, and *LINC00910* are highly conserved across primate species

It is well established that lncRNAs involved in transcriptional modulation in animals tend to exhibit low conservation across different species (7,55). We therefore decided to investigate if this remains true for the evolutionary conservation of the three lncRNAs *TILR1, TILR2*, and *LINC00910*. BLASTn analysis revealed that these lncRNAs are restricted to Old World Monkeys (OWM) and are absent in more primitive primates or other vertebrates. Remarkably, their sequence identity is unusually high for lncRNAs (Fig. 7A, Additional file 8). For example, the sequence identity of *TILR-1* and *TILR-2* between humans and chimpanzees exceeds 90%, while *LINC00910* reaches 98% identity. Even in the most distantly related OWM species, *Rhinopithecus roxellana*, sequence identity remains above 85% for *TILR-1* and *TILR-2* and 90% for *LINC00910* (Fig. 7A, Additional file 8).

**Fig 7.**
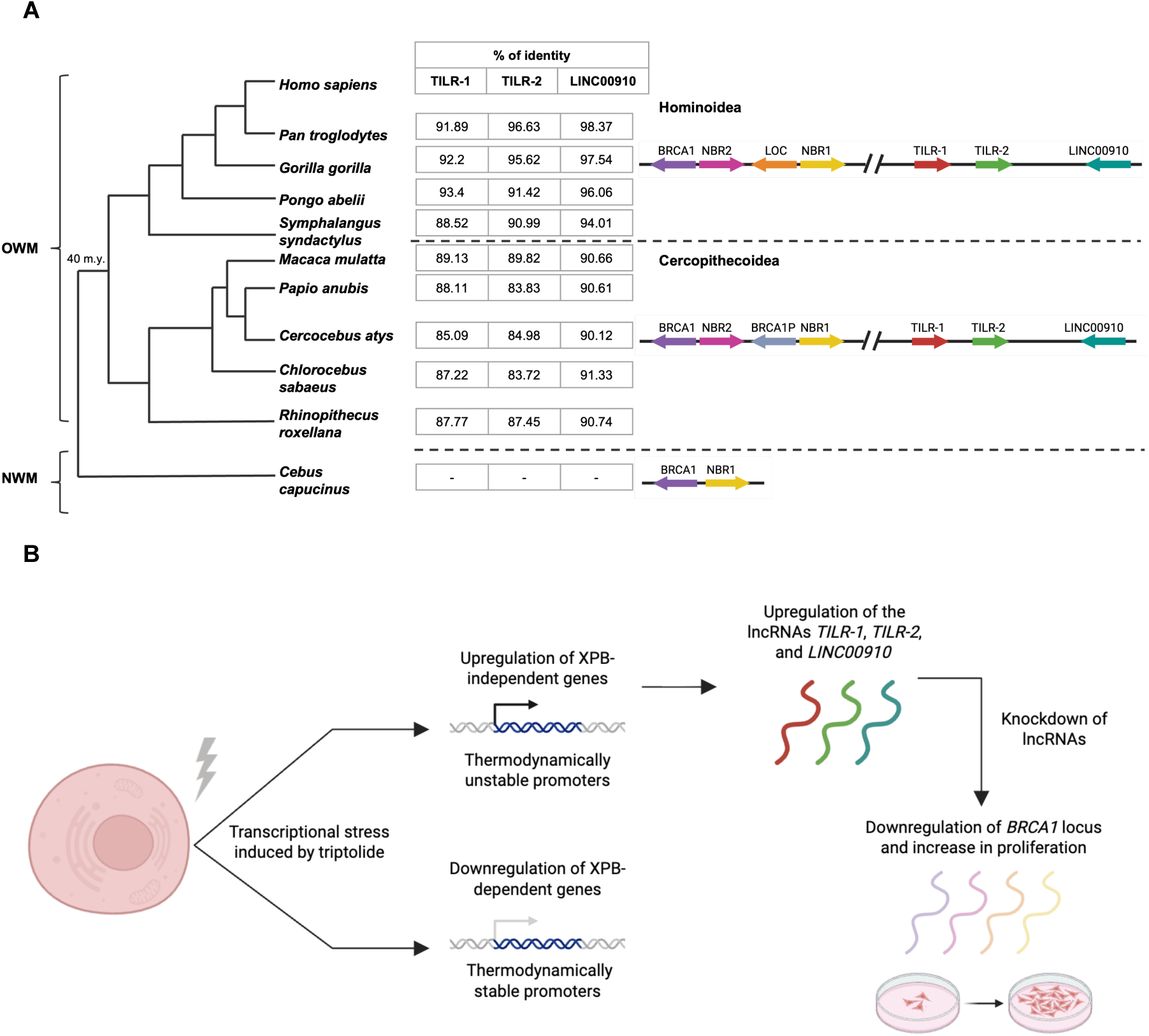
*TILR-1*, *TILR-2*, and *LINC00910* are highly conserved in primates, and are part of the transcriptional stress response induced by Triptolide treatment. **A** Conservation analysis of the *BRCA1* locus and the elements *TILR-1*, *TILR-2*, and *LINC00910* among primates. Old World Monkeys (OWM), New World Monkeys (NWM). **B** Schematic proposed model of the response of thermodynamically stable and unstable genes upon transcriptional stress induced by TPL treatment, and the upregulation of the lncRNAs *TILR-1*, *TILR-2*, and *LINC00910* in normal conditions and upon transcriptional stress.

The synteny of these three lncRNAs has been preserved since their emergence approximately 40 million years ago, highlighting their evolutionary significance (Fig. 7A). Notably, their appearance in primates coincides with a key duplication event involving *BRCA1* and *NBR1* in OWMs and with an Alu expansion (56,57). This event resulted in the formation of a four-gene locus, where *BRCA1* and *NBR2* are transcribed from one bidirectional promoter, and *LOC101929767* and *NBR1* from another (56) (Fig. 7A). These observations suggest that *TILR-1, TILR-2*, and *LINC00910* have been evolutionarily conserved to ensure proper regulation of this locus, implying that these lncRNAs co-evolved with the gene cluster and are critical for the coordinated expression of *BRCA1, NBR2, LOC101929767*, and *NBR1*.

## Discussion

Transcriptional stress response has been studied under different types of homeostasis disruptions such as heat shock, hypoxia, and oxidative stress, among others. However, the transcriptional response generated by transcriptional inhibition remains underexplored. In this study, we used TPL-treated K562 cells as a model to investigate transcriptional inhibition stress, from a genome-wide perspective, and with a particular focus on the molecular mechanisms behind the transcriptional regulation of the *BRCA1* locus carried by lncRNAs.

Previous work reported that TPL treatment, which inhibits the XPB translocase and broadly blocks transcription initiation, unexpectedly leads to the accumulation of certain transcripts as detected by RNA-seq (5). Here, we expanded our knowledge of this phenomenon by measuring nascent RNA production through PRO-seq experiments, revealing that a small subset of transcripts is actively upregulated, some of which are sufficiently stable to be detected by RNA-seq. We show that XPB-independent promoters can bypass the normal promoter melting step catalyzed by the XPB translocase because they are thermodynamically less stable than the XPB-dependent promoters. A previous report in *Drosophila* cells showed that XPB binding increases in the TSS region upon TPL treatment (35). Consistent with this we observed that this XPB accumulation occurs preferentially at XPB-dependent promoters, indicating that TPL, by blocking XPB catalytic activity, prevents its dissociation from the promoter. In contrast, promoters of XPB-independent genes do not retain XPB, suggesting that the architecture of these promoters still supports, and may even enhance, transcription initiation, even in the absence of XPB activity (Fig. 7B). Notably, this XPB independence has not been observed previously in human cells, being reported only in yeast and *in vitro* systems (31,32,34). Additionally, it has been reported that the absence of XPB does not affect transcription initiation, whereas TPL treatment does impair it in fibroblast cells, as shown by *in situ* 5EU incorporation during RNA synthesis (58). Another contributing factor may be that, under our TPL treatments, approximately 55% of RNA Pol II remains available, potentially supporting transcription at this subset of XPB-independent promoters.

By performing CUT&Tag to assess whether H3K27ac plays a role in this upregulation upon TPL treatment, we found that, under these conditions, H3K27ac does not correlate with the upregulation of this subset of transcripts. This, combined with our promoter stability analyses, suggests that promoter thermodynamic properties rather than canonical activation-associated chromatin marks underlie this transcriptional response. One possible explanation is that there is an increase in nucleosome occupancy, TFs, and co-factors at the +1 nucleosome position in these genes as a consequence of inhibiting XPB catalytic activity, which may account for the increased H3K27ac signal detected at this position. This would be consistent with the previous result in which we propose that TPL treatment prevents the dissociation of XPB, and likely of the entire PIC, from promoters (5,35). Besides this, these genes may be primed for rapid reactivation once this type of stress passes. H3K27ac modification has been reported to function as a bookmarking mark for rapid post-mitotic reset of the expression of specific genes in pluripotent stem cells (59). Interestingly, these reported genes are involved in mRNA processing and metabolism, similar to the functional enrichment that we detected in the group of hyperacetylated genes in response to TPL treatment. Although this hypothesis has not been tested under stress conditions, it is a plausible explanation for this behavior, and it would be interesting in the future to investigate these two possible scenarios.

Among the lncRNAs upregulated upon TPL treatment, we identified two novel regulatory RNAs, *TILR-1* and *TILR-2*, as well as the lncRNA *LINC00910,* all of which respond positively to this transcriptional stress. Using CRISPRi and ASO-mediated knockdowns, we demonstrated that their expression is mutually interdependent, and that this interdependence is mediated by the RNA products themselves rather than by transcription *per se*. Furthermore, these lncRNAs regulate the expression of the nearby gene cluster containing *BRCA1, NBR2, LOC101929767*, and *NBR1*. Thus, these transcripts form a coordinated lncRNA regulatory module embedded within a stress-responsive genomic region.

Loss of these transcripts results in increased cell proliferation and survival, a phenotype reminiscent of that reported previously for the *NBR2* depletion (60), and also observed in cancer cells harboring alterations in *BRCA1* or *NBR2* (61,62). *BRCA1* is a well-known tumor suppressor gene implicated in DNA repair, cell cycle regulation, and cellular growth (63,64), whereas *NBR1* is a selective autophagy receptor (65). These roles are consistent with the overall phenotype observed upon depletion of the regulatory lncRNAs, suggesting that their downregulation disrupts expression at the *BRCA1* locus and facilitates bypass of normal constraints on cellular proliferation and survival, thereby promoting uncontrolled growth under stress conditions (Fig. 7B). In the case of the lncRNA *LOC101929767*, its function remains to be elucidated.

Interestingly, the promoters within the *BRCA1* locus are sensitive to TPL, as XPB accumulates at higher levels upon treatment. However, the locus remains transcriptionally active when the associated lncRNAs are upregulated. We propose that the lncRNAs *TILR-1*, *TILR-2*, and *LINC00910* counteract the TPL sensitivity of the *BRCA1 locus* by maintaining expression of *BRCA1* locus genes. These observations support a model in which lncRNA transcriptional modules locally buffer transcriptional inhibition at sensitive genomic loci. Although the precise molecular mechanism remains to be explored, the genomic regions giving rise to these lncRNAs interact with the *BRCA1* locus. We propose that these interactions, whether direct or protein-mediated, may allow the expression of the *BRCA1* locus even when transcription is globally inhibited, potentially by maintaining the chromatin accessibility and/or by recruiting transcription factors to these promoters.

It is well established that lncRNAs are poorly conserved at the sequence level (7,55). However, there have been some examples previously described in which some lncRNAs are highly conserved, especially functional regions (66,67). Surprisingly, *TILR-1*, *TILR-2*, and *LINC00910* are highly conserved at sequence level within the OWM group of primates, and their emergence during evolution coincides with the rearrangements and duplications that occurred in the ancestral bidirectional promoter containing *BRCA1* and *NBR1* (56). During this period, Alu expansion also occurred, and these lncRNAs contain partial Alu elements. (56,57). This suggests that approximately 40 million years ago these elements emerged during Alu expansion and were positively selected, becoming conserved regulators of the rearranged *BRCA1* locus. Since we found that there are DNA-RNA interactions between the lncRNAs and the *BRCA1* locus, we propose that Alu-mediated complementarity contributes to these interactions, as complementary Alu sequences can mediate RNA–RNA interactions and enhancer–promoter specificity (68). Furthermore, it has recently been reported that a ∼40-nucleotide motif derived from Alu elements, termed FERM, is present in enhancer RNAs and is required for the modulation of gene expression mediated by the estrogen-receptor (69). Besides this, TFs bound to *TILR-1, TILR-2*, and *LINC00910* DNA regions are also bound to the two bidirectional target promoters, suggesting that these RNAs may recruit transcription factors while engaging DNA–RNA interactions, as reported for other regulatory RNAs (9,41,70–73).

Finally, we found that *TILR-1, TILR-2*, and *LINC00910* are upregulated in a variety of cell types treated with different types of transcriptional inhibitors, and notably, these RNAs are also upregulated in organoids derived from patients with pancreatic cancer treated with CK21, a TPL analog (52). This observation suggests that their function in counteracting stress could be extrapolated into systems that better recapitulate a tumor. Moreover, *LINC00910* has been previously linked to cancer (10,74,75), and in the LncSEA enrichment analysis this RNA emerges in the categories of mutations and eQTL associated with cancer. We propose that *TILR-1*, *TILR-2*, and *LINC00910* act as positive regulators of the *BRCA1* locus. Under both basal and transcriptional stress conditions, these lncRNAs contribute to the proper expression of *BRCA1* locus genes, thereby enabling an appropriate cellular stress response.

### Conclusions

Together, our findings support a model in which thermodynamically unstable promoters define a genome-wide class of XPB-independent regulatory elements that remain transcriptionally active during global transcriptional inhibition. Within this framework, stress-inducible lncRNAs such as *TILR-1*, *TILR-2*, and *LINC00910* function as components of coordinated regulatory modules that sustain transcription at sensitive genomic loci (Fig. 7B). These results reveal a previously unrecognized layer of genome organization that enables selective gene activation under transcriptional stress.

## Methods

### Cell culture

K562, MCF7, and MDA-MB-231 cell lines were originally purchased from ATCC. MCF10A-ER-Src line was donated by Dr. Kevin Struhl. K562 cells were maintained in RPMI medium (Gibco, No. 11875093), supplemented with 10% FBS (Gibco, No A5256701) and 1% streptomycin/penicillin (Invitrogen, No. 15070-063). MCF7 and MDA-MB-231 were maintained in DMEM medium (Gibco, No. 21969035) supplemented with 10% FBS and 1% streptomycin/penicillin. MCF10A-ER-Src cell line was maintained in DMEM/F12 medium (Invitrogen, No. 11039-021) supplemented with 5% Charcoal Stripped Horse Serum (Invitrogen, No. 16050-122), EGF (20 ng/ml – Peprotech, No. AF-100-15), insulin (10 μg/ml – Sigma Aldrich, No. I-1882), hydrocortisone (0.5 μg/ml – Sigma Aldrich, No. H-08888), cholera toxin (100 ng/ml – Sigma Aldrich, No. C-8052), puromycin (0.5 μg/ml – Calbiochem, No. 540411), and 1% streptomycin/penicillin. To induce the transformed phenotype of these cells, 2.5 μM Tamoxifen (Sigma Aldrich, No. T548) was added to previously plated cells for 72 hours. All cell lines were maintained at 37°C in a humidified incubator with 5% CO_2_.

### Generation of stable cell lines

K562 cell lines stably expressing dCas9 (Addgene #100091) and dCas9-KRAB (Addgene #89567) constructs were generated by nucleofecting 1×10^6^ cells with 1 μg of each plasmid respectively. Forty-eight hours after nucleofection the medium was changed with fresh RPMI medium + 0.6 mg/ml of G418 (Sigma Aldrich, No. A1720) for dCas9 cell line or 7.5 μg/ml of Blasticidin (Calbiochem, No. 3513-03-9) for dCas9-KRAB cell line. Antibiotic selection was carried out for 10 days, and monoclonal cell lines were generated by limiting dilution in 96-well plates. All monoclonal cell lines were tested by assessing the expression of dCas9 or dCas9-KRAB using flow cytometry and western blot assays with an anti-Cas9 antibody (Diagenode, C15310258). These stable cell lines were maintained with RPMI medium supplemented with 10% FBS, 1% streptomycin/penicillin, and 0.6 mg/ml G418 (K562 – dCas9) or 7.5 μg/ml Blasticidin (K562 – dCas9-KRAB).

For experiments, all cell lines were treated with DMSO (Sigma Aldrich, No. D8418) as the vehicle control or Triptolide (TPL – Tocris, No. 3253) for the specified times and concentrations.

### Flow cytometry and cellular proliferation assays

For cellular viability, apoptosis, and cell cycle measurements, 5×10^5^ K562 cells were incubated with DMSO or TPL for the specified times and concentrations according to each experiment. Treated samples were centrifugated fir 8 min at 1500 rpm and washed once with 1× PBS. 50 μl of previously diluted (1:10000 in 1× PBS) Viability Dye reagent (Invitrogen, Fixable Viability Dye eFluor 780) was added to the samples and were incubated for 30 min on ice, protected from light. Samples were centrifuged again and then incubated with 3 μl of FITC Annexin V (BioLegend, Cat. 640906) + 500 μl 1× Annexin V Binding Buffer (BioLegend, Cat. 422201) for 15 min at room temperature. Cells were washed once with 1× PBS, fixed with 100 μl of PFA at 2% for 10 min at 37°C, and permeabilized using 70% ethanol overnight at –20°C. The next day samples were washed once with 1× PBS and incubated with DAPI solution (ThermoFisher, D1306; stock 5 mg/ml diluted 1:20000 in 1× PBS) for 30 min at 37°C. Finally, samples were washed once with 1× PBS and resuspended in 200 μl of PFA 2% until analysis.

To test RNA Pol II expression, 5×10^5^ K562 cells were incubated with DMSO or TPL, centrifuged 8 min at 1500 rpm and washed once with 1 ml of 1× FACS Juice (2% FBS and 1× sodium azide in 1× PBS). Samples were permeabilized with 2 ml of methanol at –20°C for at least 1 hour. Cells were centrifuged and incubated with 20 μl of the primary antibody (8WG16, Santa Cruz Biotechnology sc-56767; dilution 1:1000 in 1× FACS Juice) for 15 min in ice, and afterward 20 μl of the secondary antibody (AlexaFluor 488 anti-mouse, dilution 1:1000 in 1× FACS Juice) were added and incubated for 45 min, for incubation time of 1 hour with both antibodies. Samples were washed with 2 ml of 1× FACS Juice and fixed with 200 μl of PFA 2%. The same procedure was followed to check the expression of dCas9 and dCas9-KRAB for the generation of stable cell lines, using the antibody against Cas9 (Diagenode, C15310258, dilution 1:400 in 1× FACS Juice).

All samples were processed on the flow cytometer BD FACSCanto II with the software BD FACSDiva and analyzed using the FlowJo v10.5.3. Three independent biological replicates were performed for each experiment, and for each replicate at least 10,000 events were counted.

To test cellular proliferation, 5×10^5^ K562 cells transfected with the corresponding antisense oligos were treated with DMSO or 250 nM TPL for 24 and 48 hours. After each time point, cells were counted three times using a Neubauer chamber and the average was calculated for each biological replicate. Five independent biological replicates were performed for each condition.

### RNA extraction for RNA-seq experiments and bioinformatic analysis

5×10^5^ K562 cells were incubated with DMSO or 250 nM of TPL for 4 hours, collected, and washed once with 1× PBS. Samples were resuspended in 200 μl of TRIzol (Invitrogen, Cat. 15596-026) and RNA was extracted following the manufacture’s protocol. 1 μg of RNA per sample was sent to Novogene for library preparation and sequencing. Libraries were enriched with oligo(dT) to capture polyadenylated transcripts, and paired-end sequencing was performed using the HiSeq PE150 platform. Three independent biological replicates were performed for each condition.

Raw data was first analyzed using FastQC software (https://www.bioinformatics.babraham.ac.uk/projects/fastqc/) to assess sample quality. Genome alignment was performed using STAR tool (76) with default parameters, using the hg19 genome as reference. Read counting was obtained with the Python package htseq-count (77), using hg19 RefSeq gene annotation. Differential expression analysis was performed with the R package DESeq2 (78), filtering by an adjusted p-value threshold of < 0.05 and a fold change of 1.5. BigWig files for IGV visualization were created using Deeptools (79).

### H3K27ac CUT&Tag experiments and bioinformatic analysis

CUT&Tag experiments were performed using the CUTANA™ CUT&Tag Kit (EpiCypher, SKU: 14-1102), following the manufacturer’s protocol. A total of 5×10^5^ K562 cells were incubated with DMSO or 250 nM TPL for 4 hours. After incubation, 1×10^5^ cells were collected for nuclei isolation and subsequent CUT&Tag processing. An overnight incubation with 3 μg of anti-H3K27ac antibody (Active Motif, No. 39133) was performed. Library amplification was carried out using 21 PCR cycles. Samples were sequenced on the HiSeq PE150 platform at Novogene. At least two independent biological replicates were performed per condition.

Bioinformatic analysis was performed following the protocol established by Zheng Y., *et al* (2020). Protocol.io. Briefly, FastQC software was first used to assess the raw data quality, and adapters were removed using Trimmomatic-0.38 (80). Alignment using the hg19 genome as reference was performed with Bowtie2 software (81), and PCR duplicates were removed with Picard 2.18.14 (http://broadinstitute.github.io/picard). To normalize the samples, the R package ChIPseqSpikeInFree (82) was used to obtain the normalization factors subsequently applied in the generation of the BigWig files and for the differential enrichment analysis. Peak calling was performed using SEACR 1.4 (83), and a peak master file with the data of all samples was created to analyze the differential enrichment analysis using the R package DESeq2 (78). Differential enriched regions were obtained filtering by an adjusted p-value threshold of < 0.05 and a fold change of 1.2. Peak annotation was performed using ChIPseeker R package (84). Metaplots were obtained using Deeptools (79).

### PRO-seq experiments and bioinformatic analysis

PRO-seq was performed following the previously described protocol (85). 10×10^6^ K562 cells were incubated with DMSO or TPL using the following concentrations and times: 1 h 250 nM, 4 h 250 nM, and 1 h 1 μM. Samples were centrifuged 5 min at 1300 rpm at 4°C, and the pellet was resuspended in 1× cold PBS. 5×10^5^ (5%) of Drosophila S2 cells were added at this point for further spike-in normalization. Cells were resuspended in 10 ml of cold permeabilization buffer (Tris-HCl, pH 7.4 10 mM, KCl 10mM, sucrose 300 mM, MgCl2 5 mM, EGTA 1 mM, IGEPAL 0.1%, Tween 20 0.05%, DTT 0.5 mM) supplemented with 1x COMPLETE EDTA-free Protease Inhibitor cocktail (Roche, Cat 11873580001), and 40 units/10 ml of SUPERaseIn RNase Inhibitor (Life Technologies, No. AM2694), and incubated on ice for 5 min. Samples were centrifuged, washed two times with cold permeabilization buffer, and resuspended in 100 μl of storage buffer (Tris-HCl 10 mM, glycerol 25%, MgCl2 5 mM, EDTA 0.1 mM, DTT 5 mM, and 1 μL of SUPERaseIn RNase Inhibitor). These permeabilized cells were flash-frozen and stored at –80°C until used.

The following run-on reaction was performed at 37°C for 5 min: 5 mM Tris-HCl pH 8.0, 150 mM KCl, 0.5% Sarkosyl, 2.5 mM MgCl2, 0.5 mM DTT, 20 μM biotin-11-CTP (Perkin Elmer, NEL542001EA), 20 μM biotin-11-UTP (Perkin Elmer, NEL543001EA), 20 μM ATP, 20 μM GTP, and 20 units/ml of SUPERaseIn RNAse Inhibitor. Total RNA was extracted using TRIzol LS (Invitrogen, No. 10296010). RNA pellet was resuspended in 20 μl of DEPC H2O and base hydrolyzed with 5 μl of 1N NaOH. Unincorporated nucleotides were removed using RNase-free P30 Columns (BioRad, No. 7326232), and biotinylated transcripts were purified three times with Dynabeads™ MyOne™ Streptavidin C1 Beads (Invitrogen, No. 65002) followed by TRIzol RNA extractions. After the first RNA purification, 3’ adaptor ligation was performed. After the second RNA purification, 5’ decapping, on-bead 5’ hydroxyl repair, and 5’ adaptor ligation was performed. Libraries were amplified using TruSeq small-RNA adaptors with 15 PCR cycles, and samples were sent for sequencing to Novogene using the HiSeq PE150 platform.

Bioinformatic analysis was performed based on previous reports (85,86). Raw data quality was first analyzed using the software FastQC (https://www.bioinformatics.babraham.ac.uk/projects/fastqc/). Adapters were removed using fastp (87) and rRNA reads were also filtered. 3’ ends were mapped using Bowtie2 (81). *Drosophila* spike-in reads were mapped against the dm6 reference genome, and human reads were mapped against the hg19 reference genome. Spike-in mapped reads were used to calculate normalization factors for each sample. BigWig files and metaplots were generated using Deeptools (79).

For gene counts, the hg19 RefSeq gene annotation was used as the transcriptome reference. Genes were divided into pause regions (TSS to +150 nt) and gene bodies (+250 nt to –250 CPS). Differential expression analysis was performed using DESeq2 (78) with the spike-in normalization factors obtained previously, filtering by an adjusted p-value threshold of < 0.05 and a fold change of 1. To classify genes into upregulated, downregulated, and highly paused, the log2 fold change values (with their respective adjusted p-value) for the pause region and gene body of each gene were plotted. Genes where the pause region and gene body are both significantly increased were called upregulated genes, whereas genes with pause region and gene body being significantly downregulated were called downregulated genes. For enhancer analysis, enhancer regions were determined using the software dREG (29). Enhancer calls from each sample were merged, and this BED file was used for enhancer counts and differential expression analysis.

### GC-content and DNA duplex free-energy analysis

DNA sequences were obtained from regions between –30 and +40 nucleotides (70 nt in total, relative to the TSS, which were corrected using GRO-cap public data). For GC-content, G and C nucleotides were counted for each sequence, and the percentage was calculated and plotted. Statistical analysis was performed using Kruskal-Wallis test in GraphPad Prism 8. For the DNA duplex free-energy analysis, the nearest-neighbor method was used following parameters from (33).

### XPB ChIP-exo and bioinformatic analysis

K562 cells were treated with DMSO or 1 μM TPL for 1 hour. Chromatin preparation and ChIP-exo were performed at the Epigenomics Facility at Cornell University – Institute of Biotechnology, using an anti-ERCC3 antibody (Cat. #HPA046077, Sigma-Aldrich).

Raw data quality was first analyzed using the FastQC software (https://www.bioinformatics.babraham.ac.uk/projects/fastqc/). Adapters were removed using fastp (87), and mapping against the hg19 reference genome was performed using Bowtie2 (81). BigWig files and metaplots were generated using Deeptools (79) and R.

### Subcellular fractionation

Subcellular fractionation was performed following the protocol described in (88). Briefly, 10×10^6^ K562 cells were incubated with DMSO or 250 nM TPL for 4 hours and then centrifuged for 5 min at 200 x g. Samples were washed with 1× PBS and divided into two tubes for RNA and protein extraction respectively. Whole-cells were resuspended in 400 μl of cell lysis buffer (10 mM Tris Ph 7.4, 150 mM NaCl, 0.15% Igepal CA-630) and incubated on ice for 5 min. Cell lysate was overlaid on top of 1 ml of Sucrose Buffer (10 mM Tris pH 7.4, 150 nM NaCl, 24% sucrose) and centrifuged at 1600 x g for 10 min. Cytoplasmic fraction was flash-frozen for the protein samples and stored at –20°C, and for RNA extraction 1 ml of TRIzol was added per 200 μl of supernatant. For nucleoplasm and chromatin separation, the pellet was first resuspended in 250 μl of glycerol buffer (20 mM Tris pH 7.4, 75 Mm NaCl, 0.5 mM EDTA, 50% glycerol) and then immediately 250 μl of Urea Nuclear Lysis Buffer (10 mM Tris pH 7.4, 1M Urea, 0.3 M NaCl, 7.5 mM MgCl2, 0.2 mM EDTA, 1% Igepal CA-630) was added. Samples were vortexed for 4 seconds and incubated on ice for 2 min. Lysate was centrifuged at 13000 x g for 2 min. Supernatant contains the nucleoplasm, which was flash-frozen for the protein samples, and for RNA extraction, 1 ml of TRIzol was added per 200 μl of the supernatant. For chromatin fraction, the pellet was resuspended in Nuclear Buffer (20 mM Hepes pH 7.6, 0.4 M NaCl, 1 mM EDTA, 1 mM EGTA) for protein extraction, and, for RNA extraction, it was resuspended in 1 ml of TRIzol.

### RNA Fluorescence In situ Hybridization (FISH)

FISH experiments for *TILR-2* and *LINC00910* were performed using a pool of 37 Quasar® 570 (*TILR-2*) and 29 CAL Fluor Red 610 (*LINC00910*) single-labeled probes (Additional file 8) designed and purchased from Stellaris Biosearch Technologies, following the manufacturer’s protocol for cells in suspension. Cells were visualized using an inverted Olympus FV1000 Confocal Microscope at 60X (2X Zoom, 1.1 NA). Images were analyzed using ImageJ software. All images shown are representative of three biological replicates, with at least 80 cells analyzed per condition.

### Western blot

For total cell protein extractions, 5×10^5^ cells were centrifuged for 5 minutes at 1300 rpm, and the pellet was resuspended in 50 μl of 1x PBS + 50 μl of 2X Laemmli Buffer (125 mM Tris pH 6.8, 4% SDS, 20% glycerol, 10% 2-mercaptoethanol, 0.004% bromophenol blue). For subcellular fractions, protein was quantified using the Bradford method, and 20 μg of protein were used for all samples. Lysates were boiled for 10 min, and proteins were separated on an SDS-PAGE gel, transferred into a nitrocellulose membrane, and blocked for 1h at RT in 5% nonfat milk dissolved in 1X TBST. Primary antibodies were incubated overnight at 4°C and secondary antibodies were incubated 1 hour at room temperature. Membranes were visualized using chemiluminescent detection with SuperSignal West Pico substrates in the ChemiDoc system (BioRad) or fluorescent detection in the Odyssey M scanner (LI-COR Biotech).

Antibodies used and their respective dilutions were:

– Lamin B1 (Abcam ab16048, 1:3000)
– GAPDH (Abcam ab9485, 1:3000)
– Histone H3 (Sigma H0164, 1:10000)
– Cas9 (Diagenode C15310258, 1:3000)
– RNA Pol II (Cell Signaling, Rpb1 NTD D864Y, 1:500)
– TUBA4A (Sigma T168, 1:500)
– Goat anti-Mouse IgG (H+L) Secondary Antibody, HRP (Invitrogen #62-6520, 1:5000)
– Goat anti-Rabbit IgG (H+L) Secondary Antibody, HRP (Invitrogen #31460, 1:5000)
– IRDye® 800CW Goat anti-Mouse IgG Secondary Antibody (LICOR #926-32210, 1:10000)
– IRDye® 800CW Goat anti-Rabbit IgG Secondary Antibody (LICOR # 926-32211, 1:10000)

### RNA extraction and RT-qPCR

Total RNA extractions were performed using TRIzol and according to manufacturer’s protocol. Between 300 ng and 1 μg of RNA was used for reverse transcription using M-MLV reverse transcriptase (Invitrogen, Cat. 28025-013) with oligo dT and random hexamer primers (unless otherwise specified in the text). For qPCR experiments, SsoAdvanced Universal SYBR Green Supermix (BioRad, No. 1725271) or SYBR Green I (Invitrogen, S7567) was used, and reactions were performed in the CFX Opus Real-Time PCR system or Roche LightCycler 480 machine. For each sample, three technical replicates were performed, and the average of these replicates represented the value for the respective biological replicate. At least three biological replicates were analyzed for each experiment.

For relative quantification, data was analyzed using the 2^-ΔCt^ method, in which the ΔCt = Ct of target gene – Ct of reference gene (GAPDH). For absolute quantification, DNA corresponding to *TILR-1*, *TILR-2*, and *LINC00910* regions were cloned into pBlueScript plasmid, and these constructs were used to create a standard curve with five serial dilutions of 1:5, starting with 10 ng of material. In all cases, the R value was > 0.99. Using these curves and the Ct values obtained from the qPCR experiments, concentration values were extrapolated. One-way ANOVA tests were performed in all cases to assess statistical significance using the software GraphPad Prism 8. Primer sequences for RT-qPCRs are found in Additional file 9.

### ASOs design and transfection

Antisense oligos (ASOs) were designed based on previously published protocols (10,45). Briefly, ASOs were synthesized as RNase H-recruiting locked nucleic acid (LNA) phosphorothioate gapmers (Affinity Plus^TM^, IDT), with 10 DNA nucleotides flanked by 3 LNA nucleotides at the 5’ and 3’ ends. For the design, the sequences were analyzed using the RNAfold WebServer, University of Vienna (89), and from the predicted secondary structure, a 16-nucleotide region was chosen based on the low probability of internal base pairing and high specificity. As a control, an ASO against GFP was used. ASO sequences are found in Additional file 9.

50 nM of each ASO or the pool of ASOs were nucleofected into K562 cells. Six hours after nucleofection, RNA extraction was performed using TRIzol, and knockdown efficiency was tested by RT-qPCR.

### sgRNA constructs and CRISPRi/a experiments

For CRISPRi/a experiments, sgRNAs were designed using Benchling software (https://benchling.com). These were cloned individually into the pSPgRNA plasmid (Addgene #47108), following the protocol described in https://media.addgene.org/cms/filer_public/6d/d8/6dd83407-3b07-47db-8adb-4fada30bde8a/zhang-lab-general-cloning-protocol-target-sequencing_1.pdf. All plasmids were corroborated by Sanger sequencing.

For CRISPRi experiments, 1×10^6^ cells were nucleofected with 1 μg of each sgRNA into K562-dCas9-KRAB or K562-dCas9 stable cell lines. 48 hours after nucleofection, 5×10^5^ cells were treated with DMSO or 250 nM TPL for 4 hours, and silencing efficiency was measured by extracting RNA using TRIzol followed by RT-qPCR. As a negative control, cell lines were transfected with 1 μg of the empty pSPgRNA plasmid.

For CRISPRa experiments, 1×10^6^ cells were nucleofected with 1 μg of each sgRNA + 3 μg of dCas9-VPR plasmid (Addgene #63798). Overexpression was measured 48 hours after nucleofection by extracting RNA using TRIzol followed by RT-qPCR. As a negative control, cell lines were transfected with 1 μg of the empty pSPgRNA plasmid. sgRNAs sequences are found in Additional file 9.

### Analysis of public RNA-seq, ChIP-seq, Hi-C, and ChAR-seq datasets

The GEO numbers corresponding to the analyzed data are listed in Additional file 7. RNA-seq and ChIP-seq raw data were downloaded from the GEO Database or ENCODE. RNA-seq data were analyzed as previously described in the Methods section. For ChIP-seq analysis, raw data were mapped to the hg19 reference genome using Bowtie2 software (81). Peak calling was performed using MACS3 (90), and BigWig files were created using Deeptools (79).

For virtual 4C visualization, we downloaded public Hi-C data performed in K562 cells and produced a BEDPE file using Juicer 1.6 (91). For ChAR-seq data, we downloaded the raw data from the GEO Database and analyzed it following previously published pipelines (53,92), with a prior filtering step in which we enriched for *TILR-1*, *TILR-2*, and *LINC00910* in the original FASTQ file using BLASTn before splitting the RNA–DNA chimeras and aligning them to the genome.

### Conservation analysis

Orthologs of *TILR-1*, *TILR-2*, and *LINC00910* were searched using BLASTn. FASTA files were queried against all vertebrates, and the results with >50% of identity and >50% query were retained, based on previous work (93). Accession numbers for these sequences are found in Additional file 8.

## Declarations

### Ethics approval and consent to participate

Not applicable.

### Consent for publication

Not applicable.

### Availability of data and materials

The raw and processed data files generated in this study are available in NCBI GEO database with the following accession numbers: GSE299949 (PRO-seq), GSE299950 (ChIP-exo), GSE299951 (CUT&Tag), and GSE299952 (RNA-seq).

### Authors’ contributions

Conceptualization: S.C.-R., J.T.L., and M.Z.; Investigation: S.C.-R.; Methodology: S.C.-R. and R.V. Formal analysis: S.C.-R.; Validation: S.C.-R.; Resources and supervision: M.F.M., J.T.L., and M.Z. Writing – original draft: S.C.-R. and M.Z.; Writing – review and editing: all the authors. Funding acquisition: J.T.L. and M.Z.

## Acknowledgements

We thank all the members of the Zurita and Lis lab for helpful discussions and ideas. We also thank Martha Vazquez for suggestions and ideas. We thank Andrea Ortega for technical assistance, and Erika Melchy-Pérez for technical assistance and use of the flow cytometer. We also thank Jerome Verleyen for technical support and access to HPC infrastructure at the Unidad Universitaria de Secuenciación Masiva y Bioinformática, Instituto de Biotecnología (UNAM), which is part of the Laboratorio Nacional de Apoyo Tecnológico a las Ciencias Genómicas (CONAHCyT). We thank Andres Saralegui and the National Laboratory of Advance Microscopy at the Instituto de Biotecnología-UNAM for technical advice in the use of the confocal microscopy. We also thank the Epigenomics Facility (RRID:SCR_021287) at the Cornell Institute of Biotechnology for ChIP-exo experiments. S.C.– R. was supported by a Fulbright García-Robles fellowship and a scholarship from CONAHCyT (957622) as a student of the Programa de Doctorado en Ciencias Bioquímicas at the Universidad Nacional Autónoma de México.

## Additional File 1: Supplementary Figures S1-S12

**Fig S1.**
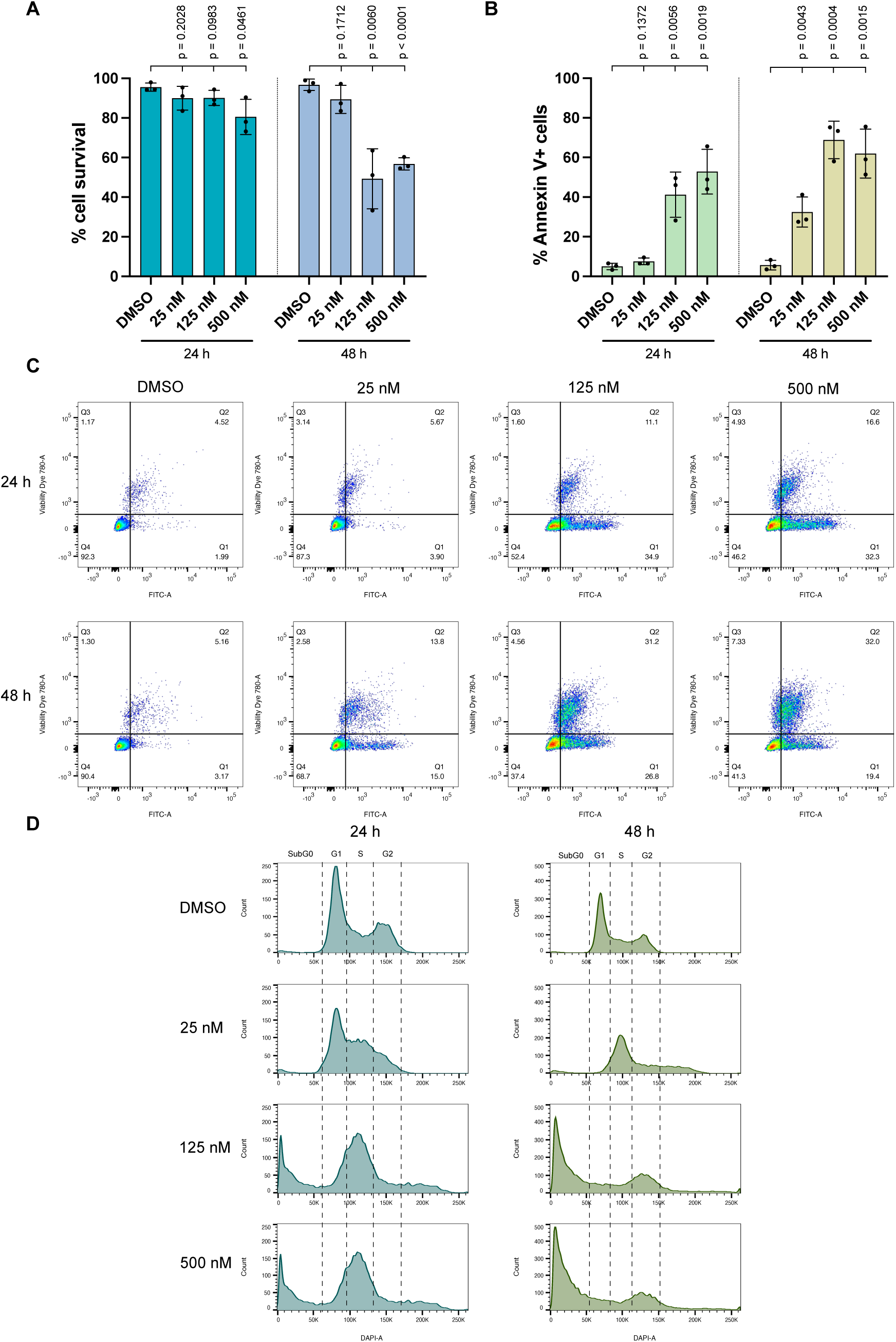
Triptolide treatment for 24 and 48 hours induces apoptosis-mediated cell death in K562 cells. **A** Cell survival and **B** apoptosis (measured by Annexin V positive cells) analysis upon triptolide (TPL) treatment at concentrations ranging from 25 nM to 500 nM for 24 and 48 hours. Data represent mean ± SD of n = 3 biological replicates. Exact p-value is indicated above (one-way ANOVA test). **C** Comparative plot showing cell survival (x axis) versus apoptosis (y axis – measured by Annexin V staining) under different TPL treatments for 24 and 48 hours. Plots are a representative example from three biological replicates. **D** Cell cycle assay in cells treated with TPL, measured by DAPI staining. Plots are a representative example from three biological replicates.

**Fig S2.**
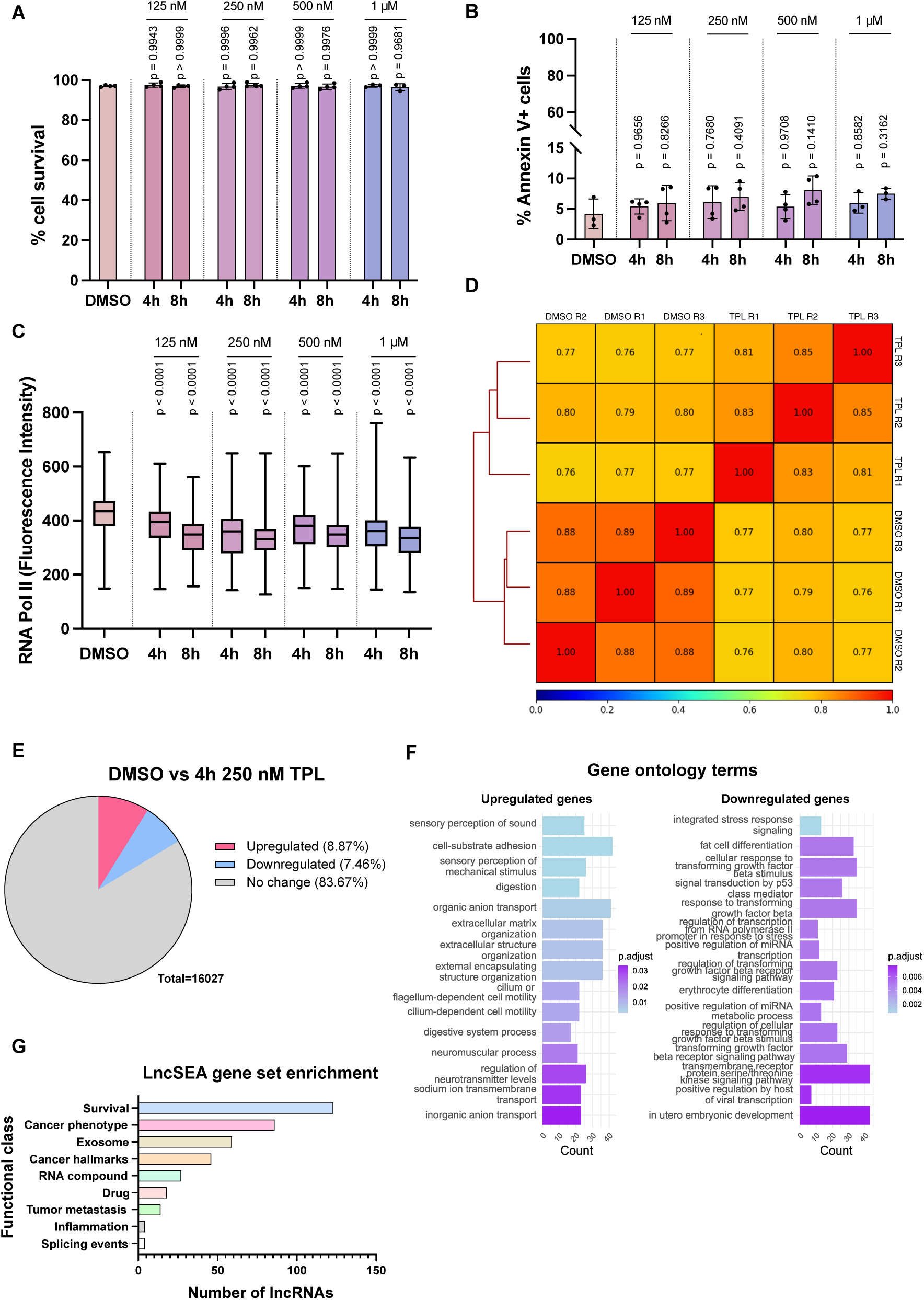
Triptolide treatments for 4 and 8 hours do not affect cell viability but cause a downregulation in RNA Pol II protein expression and in global RNA expression. **A** Cell survival and **B** apoptosis (measured by Annexin V positive cells) upon Triptolide (TPL) treatment at concentrations ranging from 125 nM to 1 μM for 4 and 8 hours. Data represent mean ± SD of n = 3 biological replicates. Exact p-value is indicated above (one-way ANOVA test). **C** Quantification of total RNA Pol II by flow cytometry in K562 cells treated with TPL for 4– and 8-hours at concentrations ranging from 125 nM to 1 μM. Data represents mean ± SD of fluorescence intensity in each cell; n = 3 biological replicates, 10000 cells per biological replicate. Exact p-value is indicated above (one-way ANOVA test). **D** Correlation heatmap between RNA-seq samples, calculated by Pearson test. **E** Percentages of upregulated, downregulated, and unchanged transcripts in cells treated with 250 nM for 4 hours, according to the RNA-seq data. **F** Gene ontology plots of the upregulated and downregulated genes found in the RNA-seq data. **G** LncRNAs functional gene enrichment of the upregulated lncRNAs found in the RNA-seq data using the software LncSEA 2.0.

**Fig S3.**
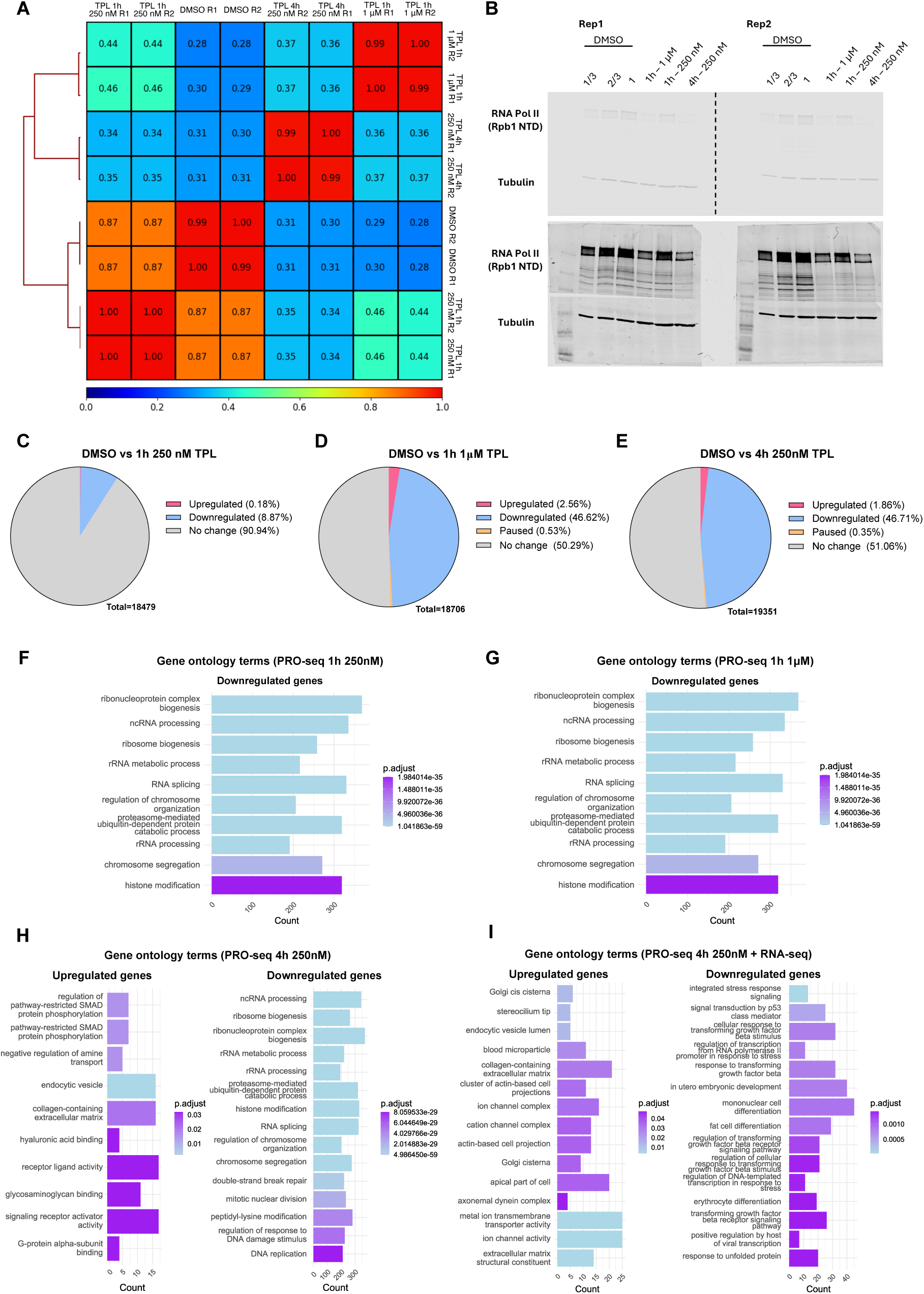
PRO-seq experiments showed a transcriptional stress response in K562 upon a variety of TPL treatments. **A** Correlation heatmap between PRO-seq samples, calculated by Pearson test. **B** Western blot showing the downregulation of total RNA Pol II upon Triptolide treatments. Tubulin was used as loading control, and two biological replicates were performed. Upper panel: less exposed Western blot membrane. Lower panel: more exposed Western blot membrane. **C-E** Percentages of upregulated, downregulated, highly paused, and unchanged genes in cells treated with **C** 1h 250 nM, **D** 1h 1 μM, and **E** 4h 250 nM, according to the PRO-seq data. **F-I** Gene ontology plots of the upregulated and downregulated genes found in the PRO-seq and RNA-seq data: **F** downregulated genes upon 1h 250 nM, **G** downregulated genes upon 1h 1 μM, **H** upregulated and downregulated genes upon 4h 250 nM, **I** and upregulated and downregulated genes upon 4h 250 nM PRO-seq + RNA-seq.

**Fig S4.**
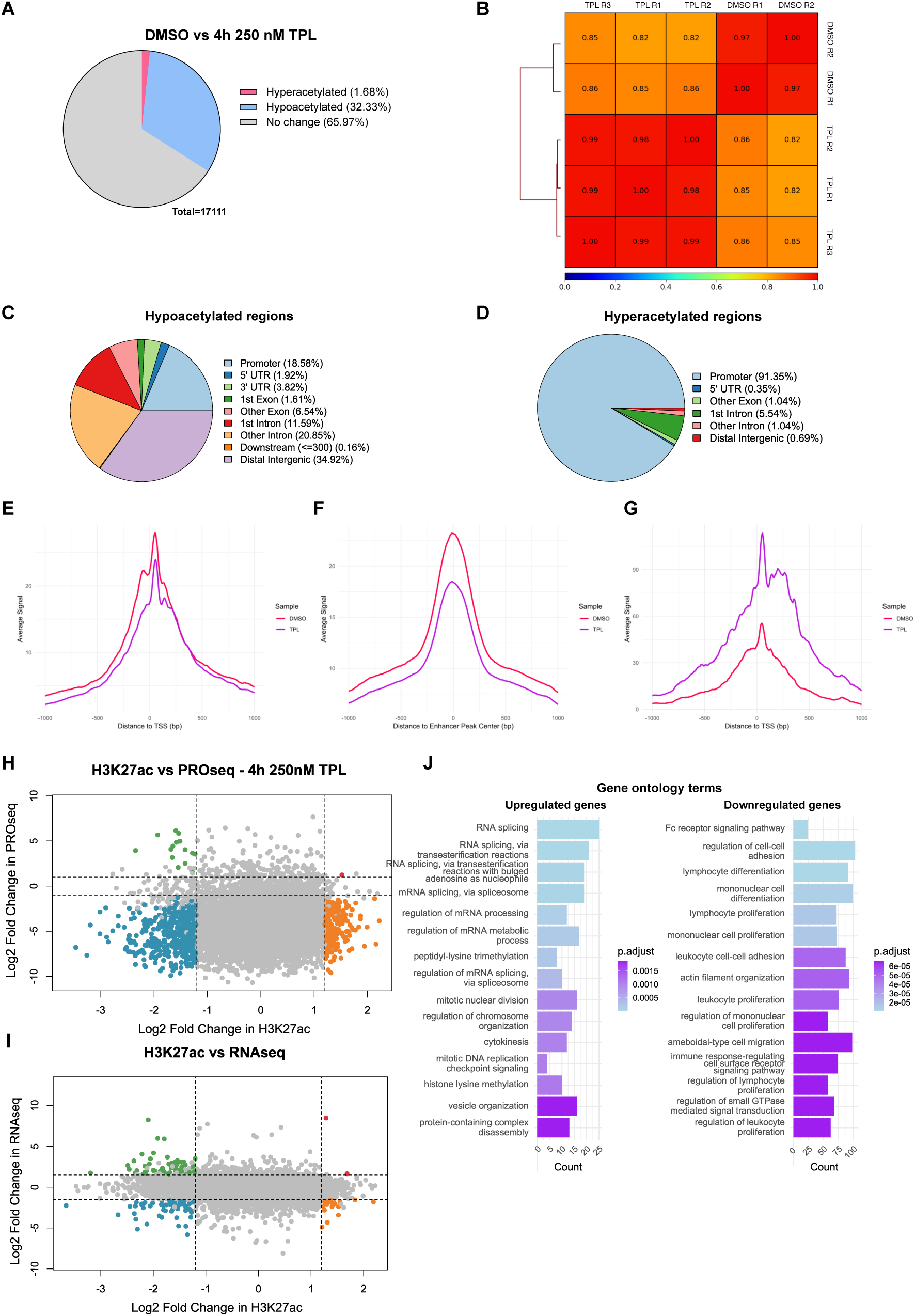
CUT&Tag H3K27ac does not correlate with the upregulation of the genes upon TPL treatment. **A** Percentages of hyperacetylated, hypoacetylated, and unchanged acetylated regions in cells treated with 250 nM for 4 hours. n = 3 biological replicates per condition were performed. **B** Correlation heatmap between CUT&Tag samples, calculated by Pearson test. **C-D** Genomic distribution of the mark H3K27ac in **C** downregulated or **D** upregulated acetylated regions after TPL treatment. **E-G** Metagene analysis of the **E** promoters and **F** enhancers that showed downregulated acetylation, as well as the promoters that presented **G** upregulated acetylation upon TPL treatment. Data is shown as an average signal in 10-nt bins. **H-I** Comparison between H3K27ac CUT&Tag (x-axis) with **H** PRO-seq 4h 250nM (y-axis) or **I** RNA-seq data (y-axis). Red dots = upregulated genes + higher enrichment of H3K27ac in the promoters; blue dots = downregulated genes + lower levels of acetylation; orange dots = downregulated genes with higher levels of acetylation; green dots = upregulated genes with lower levels of acetylation. **J** Gene ontology plots of the hyperacetylated and hypoacetylated genes found in the CUT&Tag data.

**Fig S5.**
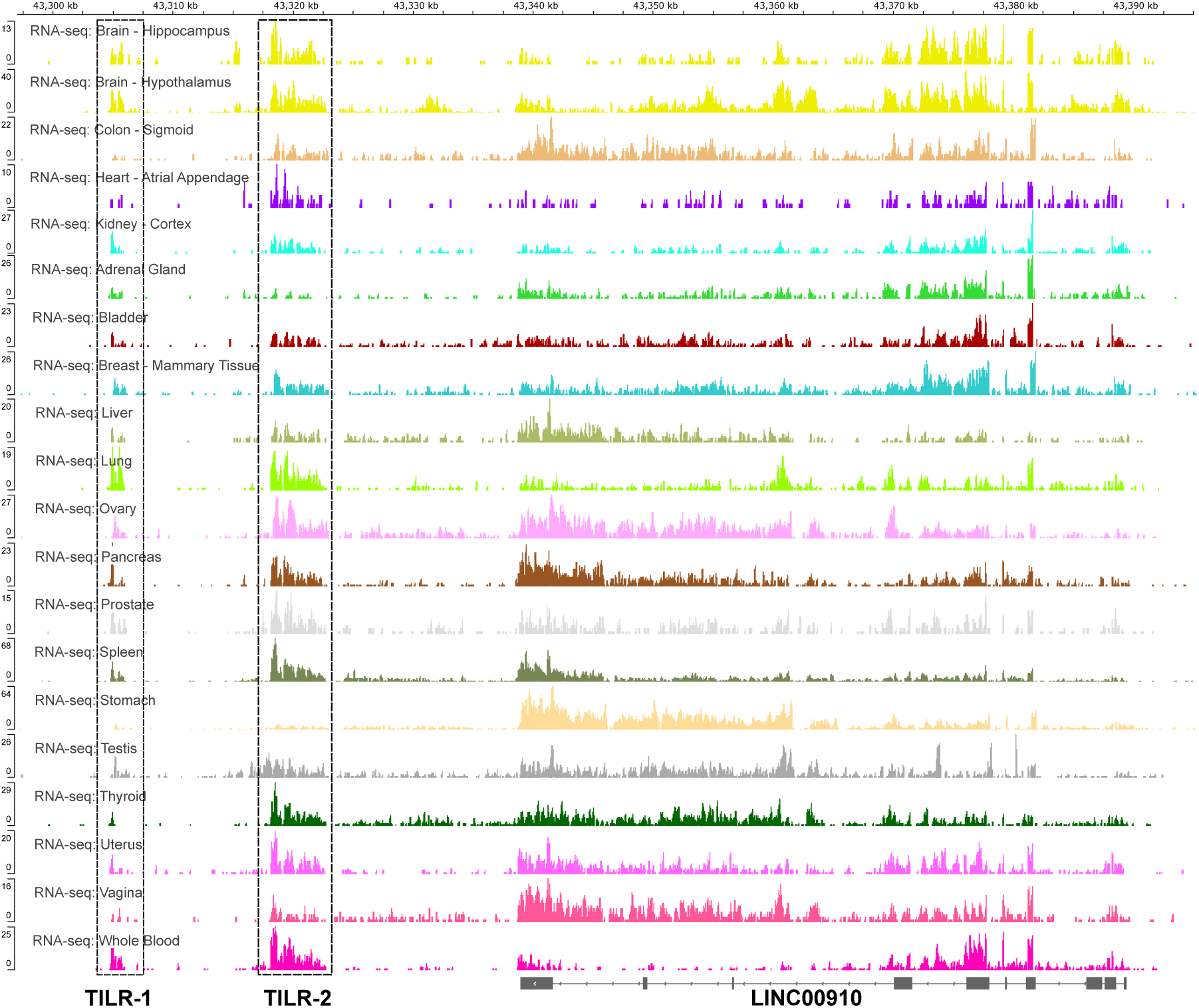
LncRNAs *TILR-1*, *TILR-2*, and *LINC00910* are expressed across human tissues. Data was taken from GTEX portal.

**Fig S6.**
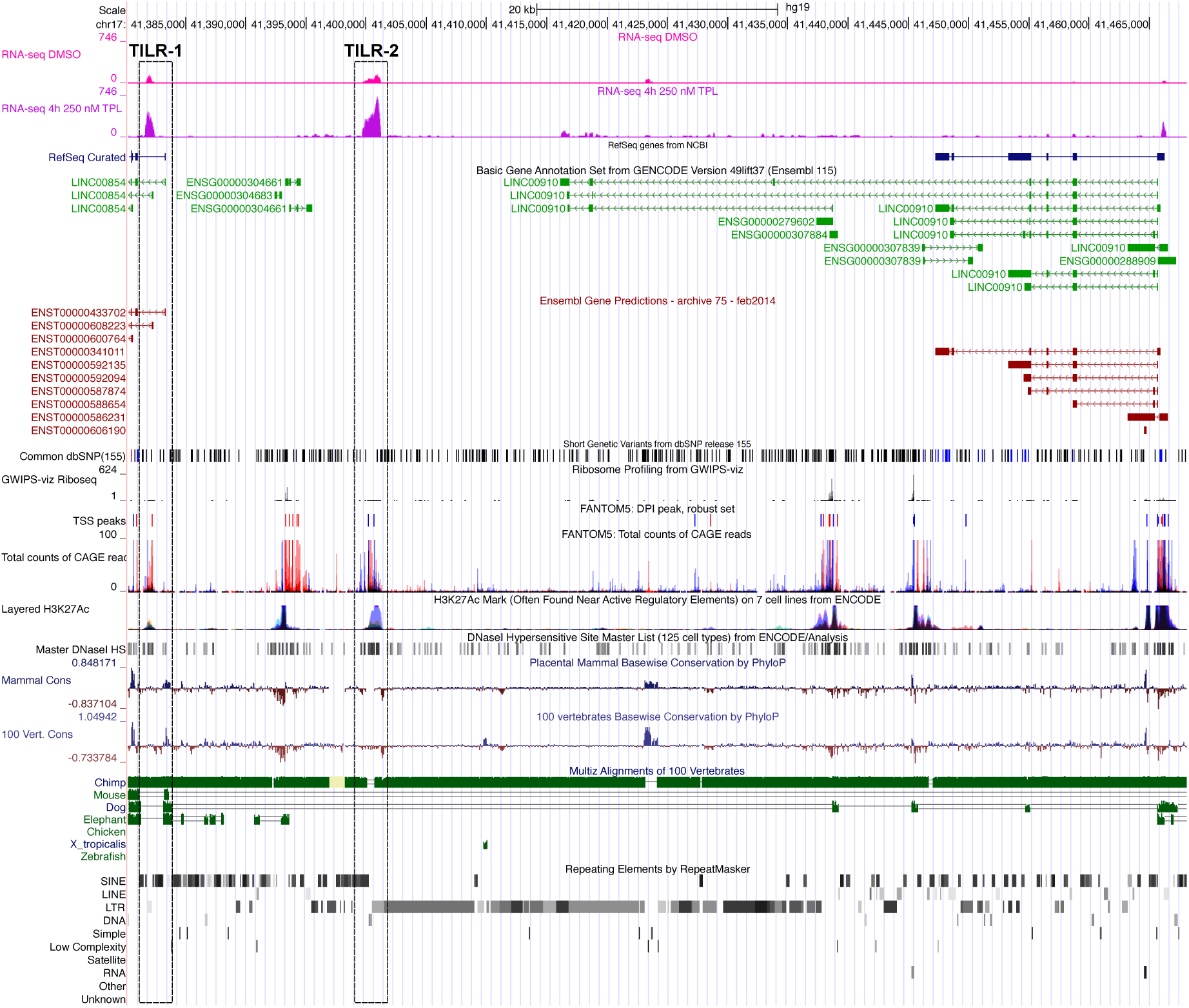
The discovery of the two novel lncRNAs *TILR-1* and *TILR-1* in chromosome 17q31. UCSC Genome Browser image of the region corresponding to *TILR-1*, *TILR-2*, and *LINC00910*. Shown are data from Ensembl annotation, dbSNPs, Ribo-seq, CAGE and DNase experiments, as well as conservation analysis in mammals and vertebrates, and the repeating elements across this region.

**Fig S7.**
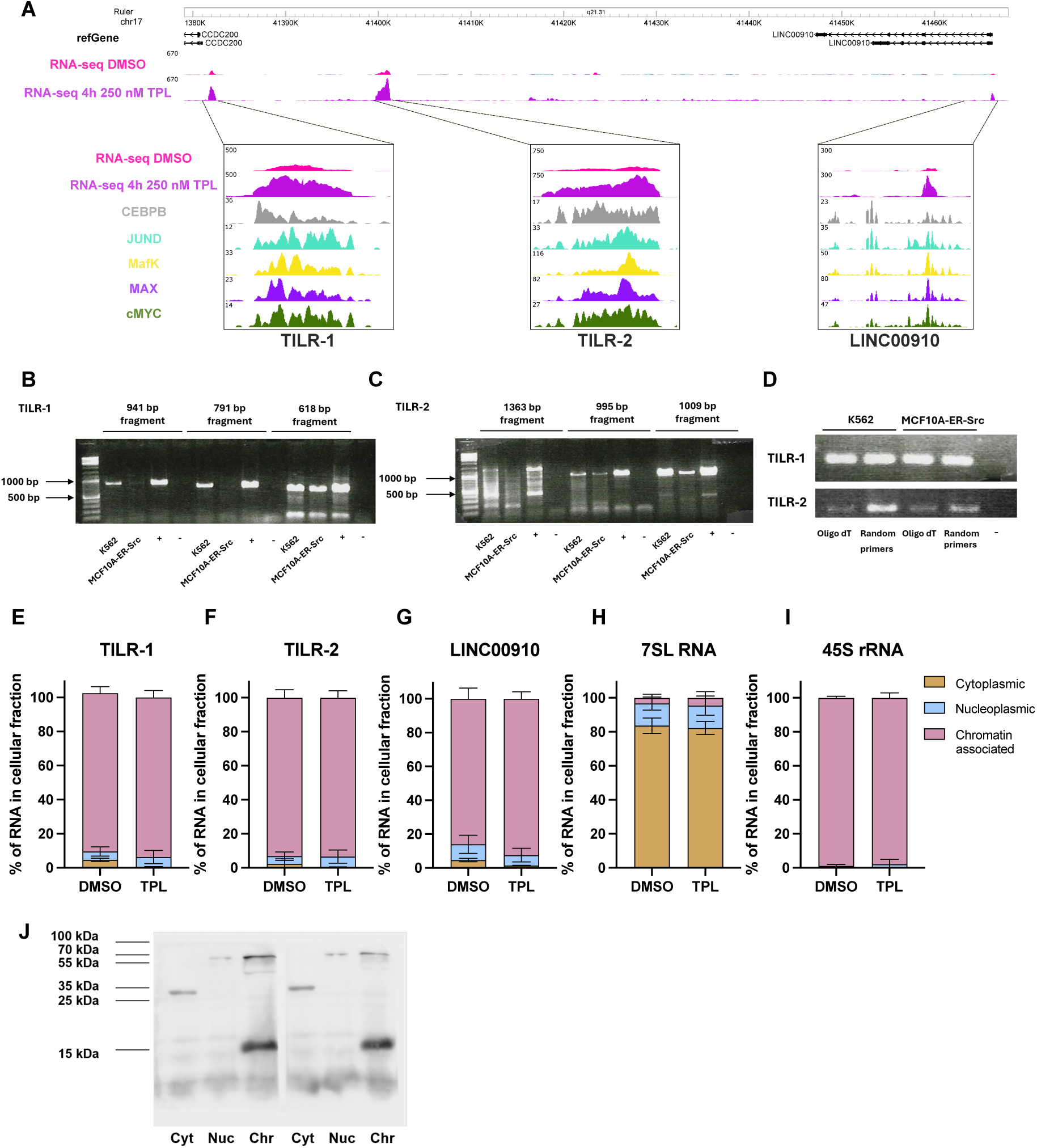
LncRNAs *TILR-1*, *TILR-2*, and *LINC00910* have characteristics of regulatory RNAs. **A** Genome browser image of the region corresponding to *TILR-1*, *TILR-2*, and *LINC00910*. Shown are RNA-seq data in K562 cells treated with DMSO or 4h 250 nM TPL, along with the ENCODE data of the transcription factors CEBPB, JUND, MafK, MAX, and c-Myc. **B-C** RT-PCR showing the length of **B** TILR-1 and **C** TILR-2 lncRNAs in K562 and MCF10A-ER-Src cells. + = positive control (genomic DNA); – = negative control (water). **D** RT-PCR to determine the poly-A presence on the TILR-1 and TILR-2 transcripts in K562 and MCF10A-ER-Src cells. **E-I** Cellular fractionation experiments followed by RT-qPCRs evaluating the expression of *TILR-1* **E**, *TILR-2* **F**, *LINC00910* **G**, 7SL RNA (cytoplasmic marker) **H** and 45S rRNA precursor (chromatin marker) **I** in K562 cells. **J** Western blot corroborating the correct cellular fractionation, using the following markers for each fraction: Cytoplasm (Cyt): GAPDH – 37 kDa; Nucleoplasm (Nuc): Laminin-B – 66 kDa; Chromatin (Chr): Histone H3 – 17 kDa. Two biological replicates are shown.

**Fig S8.**
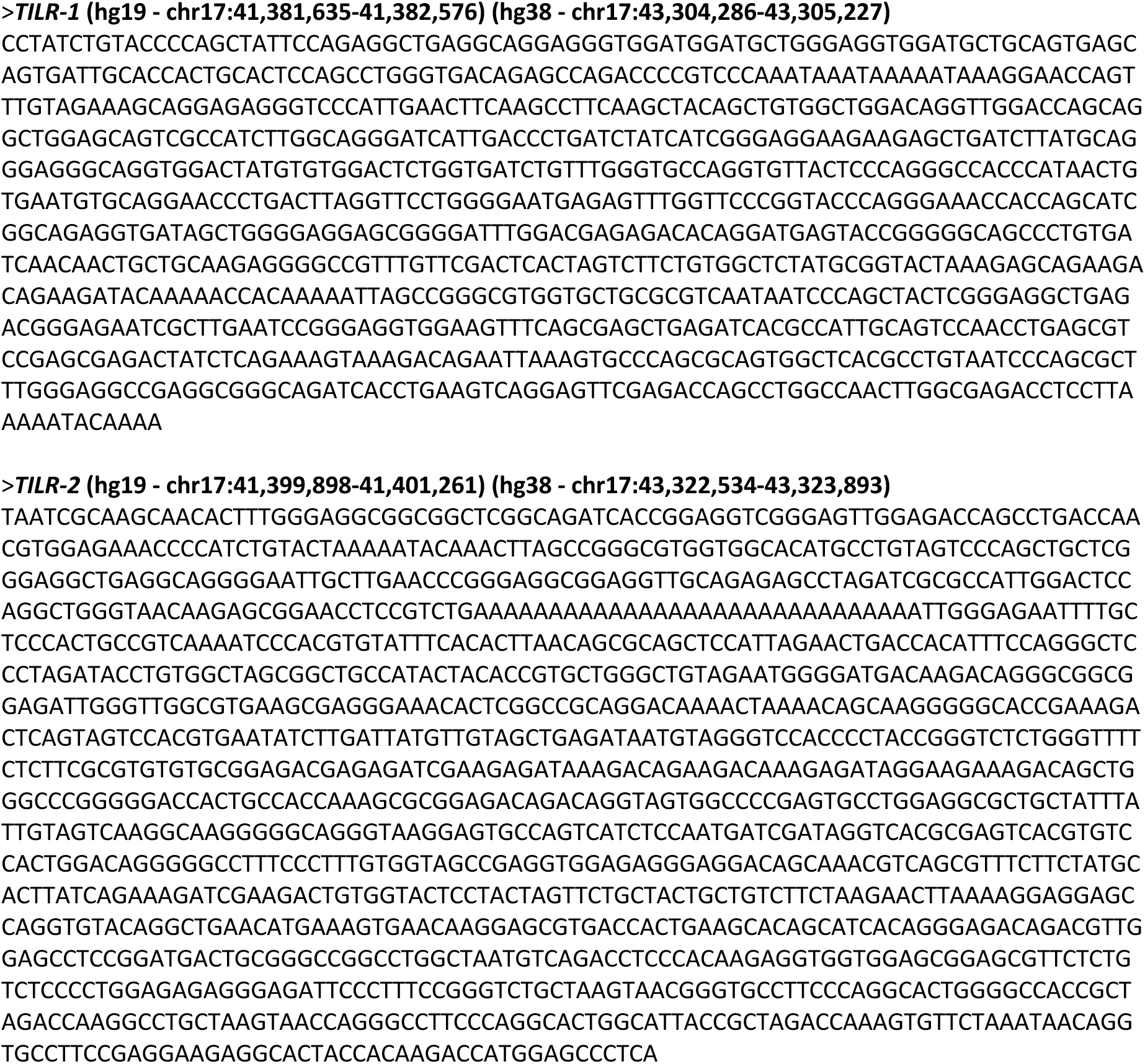
Complete sequence and coordinates corresponding to the lncRNAs *TILR-1* and *TILR-2*.

**Fig S9.**
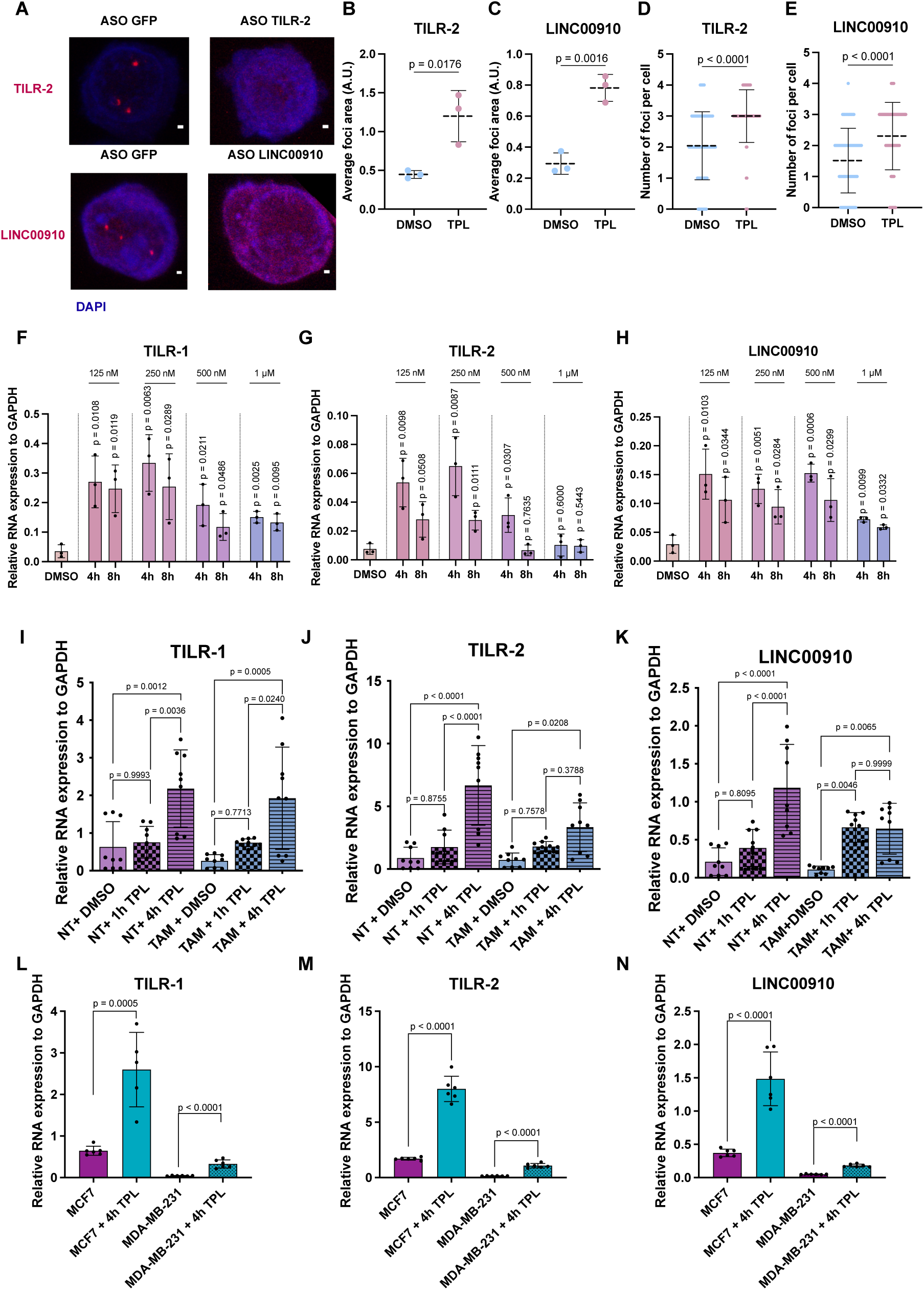
LncRNAs *TILR-1*, *TILR-2*, and *LINC00910* are upregulated in different cell lines upon Triptolide treatment. **A** Representative images of *TILR-2* and *LINC00910* FISH in K562 cells transfected with ASO GFP (control) or ASO *TILR-2* and ASO *LINC00910*, respectively. Scale bar = 1 μm. **B-C** Dot plot showing the quantification of **B** *TILR-2* and **C** *LINC00910* FISH foci area in individual cells (A.U. = arbitrary units). Each dot represents the average quantification of cells in each replicate. Exact p-value is indicated above (Mann-Whitney test). **D-E**. Dot plot showing the number of **D** *TILR-2* and **E** *LINC00910* foci per cell. Exact p-value is indicated above (Mann-Whitney test). **F-H** RT-qPCRs evaluating the expression of **F** *TILR-1*, **G** *TILR-2*, **H** and *LINC00910* in K562 cells treated with DMSO or TPL at different concentrations (125 nM to 1 μM) for 4 or 8 hours. Data represent mean ± SD of at least n = 3 biological replicates. Exact p-value is indicated above (one-way ANOVA test). **I-K** RT-qPCR to measure expression of **I** *TILR-1*, **J** *TILR-2*, and **K** *LINC00910* in MCF10A-ER-Src cells non-transformed (NT) and transformed (TAM) treated with DMSO or TPL 125 nM for 1 and 4 hours. Data represent mean ± SD of at least n = 3 biological replicates. Exact p-value is indicated above (one-way ANOVA test). (**L-N**) RT-qPCR to measure expression of **L** *TILR-1*, **M** *TILR-2*, **N** and *LINC00910* in MCF7 and MDA-MB-231 treated with DMSO or TPL 125 nM for 4 hours. Data represent mean ± SD of at least n = 3 biological replicates. Exact p-value is indicated above (one-way ANOVA test).

**Fig S10.**
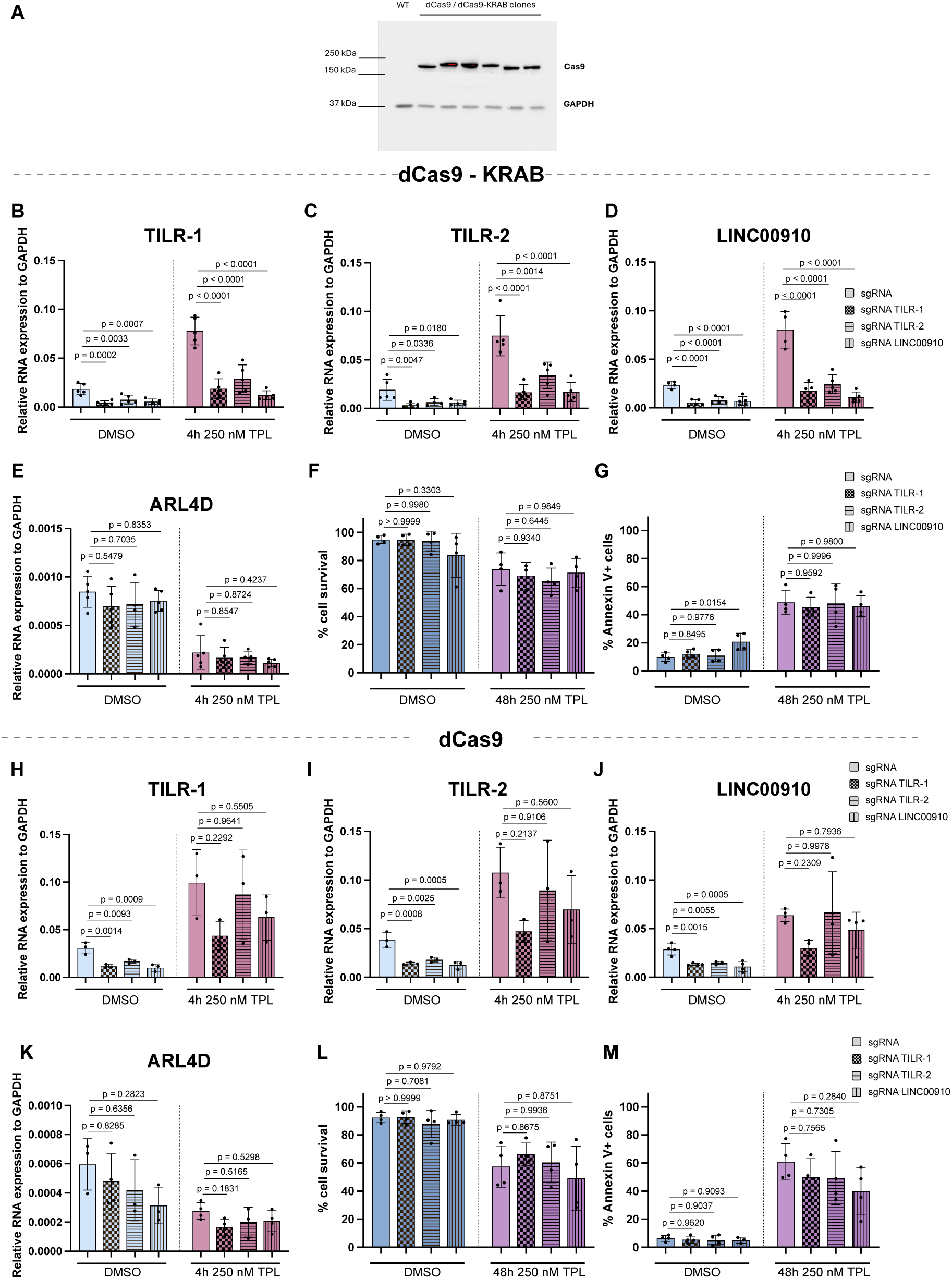
LncRNAs *TILR-1*, *TILR-2*, and *LINC00910* expression is interdependent. **A** Western blot showing the generation of stable cell lines expressing dCas9-KRAB and dCas9 constitutively. **B-E** RT-qPCR to measure expression of **B** *TILR-1*, **C** *TILR-2*, **D** *LINC00910*, and **E** *ARL4D* 48 hours after nucleofection of sgRNAs targeting these RNAs for interference using the K562-dCas9-KRAB stable cell line + 4 hours of treatment with DMSO or 250 nM TPL. As negative control, sgRNA plasmid without gRNAs was used. Data represent mean ± SD of n = 5 biological replicates. Exact p-value is indicated above (one-way ANOVA test). **F** Cell survival and **G** apoptosis measured by Annexin V positive cells analysis 48 hours after nucleofection of sgRNAs targeting separately the RNAs *TILR-1*, *TILR-2*, and *LINC00910* for silencing using the cell line K562-dCas9-KRAB + 48 hours of treatment with DMSO or 250 nM TPL. Data represent mean ± SD of n = 4 biological replicates. Exact p-value is indicated above (one-way ANOVA test). **H-K** RT-qPCR to measure expression of **H** *TILR-1*, **I** *TILR-2*, **J** *LINC00910*, and **K** *ARL4D* 48 hours after nucleofection of sgRNAs targeting these RNAs for interference using the K562-dCas9 stable cell line + 4 hours of treatment with DMSO or 250 nM TPL. As negative control, sgRNA plasmid without gRNAs was used. Data represent mean ± SD of n = 3 biological replicates. Exact p-value is indicated above (one-way ANOVA test). n.s., not significant. **L** Cell survival and **M** apoptosis measured by Annexin V positive cells analysis 48 hours after nucleofection of sgRNAs targeting separately the RNAs TILR-1, TILR-2, and LINC00910 for silencing using the cell line K562-dCas9 + 48 hours of treatment with DMSO or 250 nM TPL. Data represent mean ± SD of n = 3 biological replicates. Exact p-value is indicated above (one-way ANOVA test).

**Figure S11.**
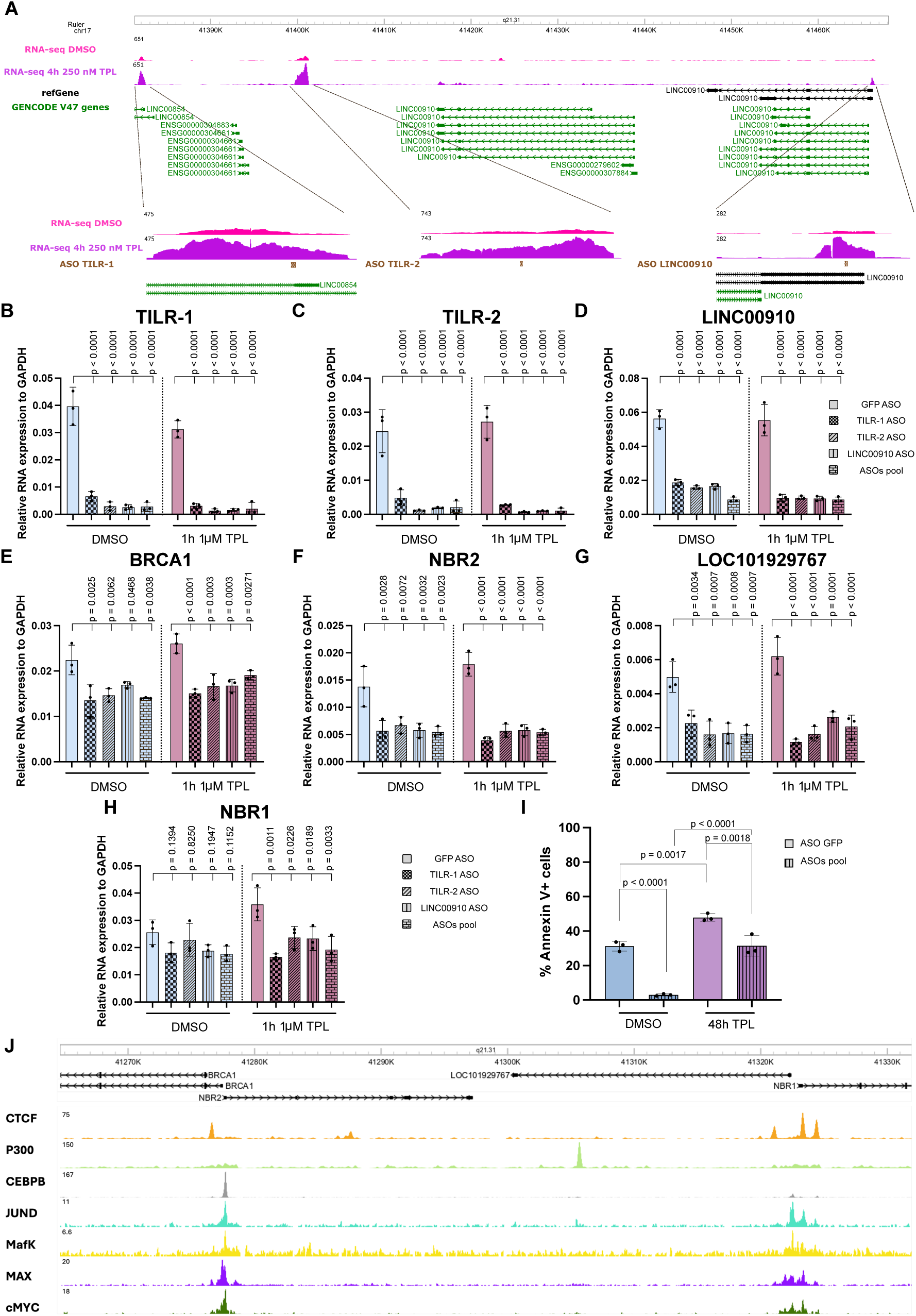
LncRNAs *TILR-1*, *TILR-2*, and *LINC00910* regulate the expression of *BRCA1*, *NBR1*, *NBR2*, and *LOC101929767*. (**A**) Genome browser image of the region corresponding to *TILR-1*, *TILR-2*, and *LINC00910*. Shown are RNA-seq data from K562 cells treated with DMSO or with 250 nM TPL for 4 hours, along with the sequences of each ASO used against each RNA, shown in brown. (**B-H**) RT-qPCR to measure expression of (**B**) *TILR-1*, (**C**) *TILR-2*, (**D**) *LINC00910*, (**E**) *BRCA1*, (**F**) *NBR2*, (**G**) *LOC101929767*, and (**H**) *NBR1* 6 hours after nucleofection of ASOs for the knockdown of the lncRNAs *TILR-1*, *TILR-2*, and *LINC00910* + 1 hour of treatment with DMSO or 1 μM TPL. As negative control, an ASO against GFP was used. Data represent mean ± SD of n = 3 biological replicates. Exact p-value is indicated above (one-way ANOVA test). (**I**) Apoptosis measured by Annexin V positive cells analysis of transfected cells with a pool of ASOs against *TILR-1*, *TILR-2*, and *LINC00910*, or against GFP as a control. These cells were treated with DMSO or 250 nM TPL for 48 hours. Data represent mean ± SD of n = 3 biological replicates. Exact p-value is indicated above (one-way ANOVA test). (**J**) Genome Browser image corresponding to the *BRCA1* locus. ChIP-seq data of the transcriptional factors CTCF, P300, CEBPB, JUND, MafK, MAX, and c-Myc was downloaded from ENCODE.

**Fig S12.**
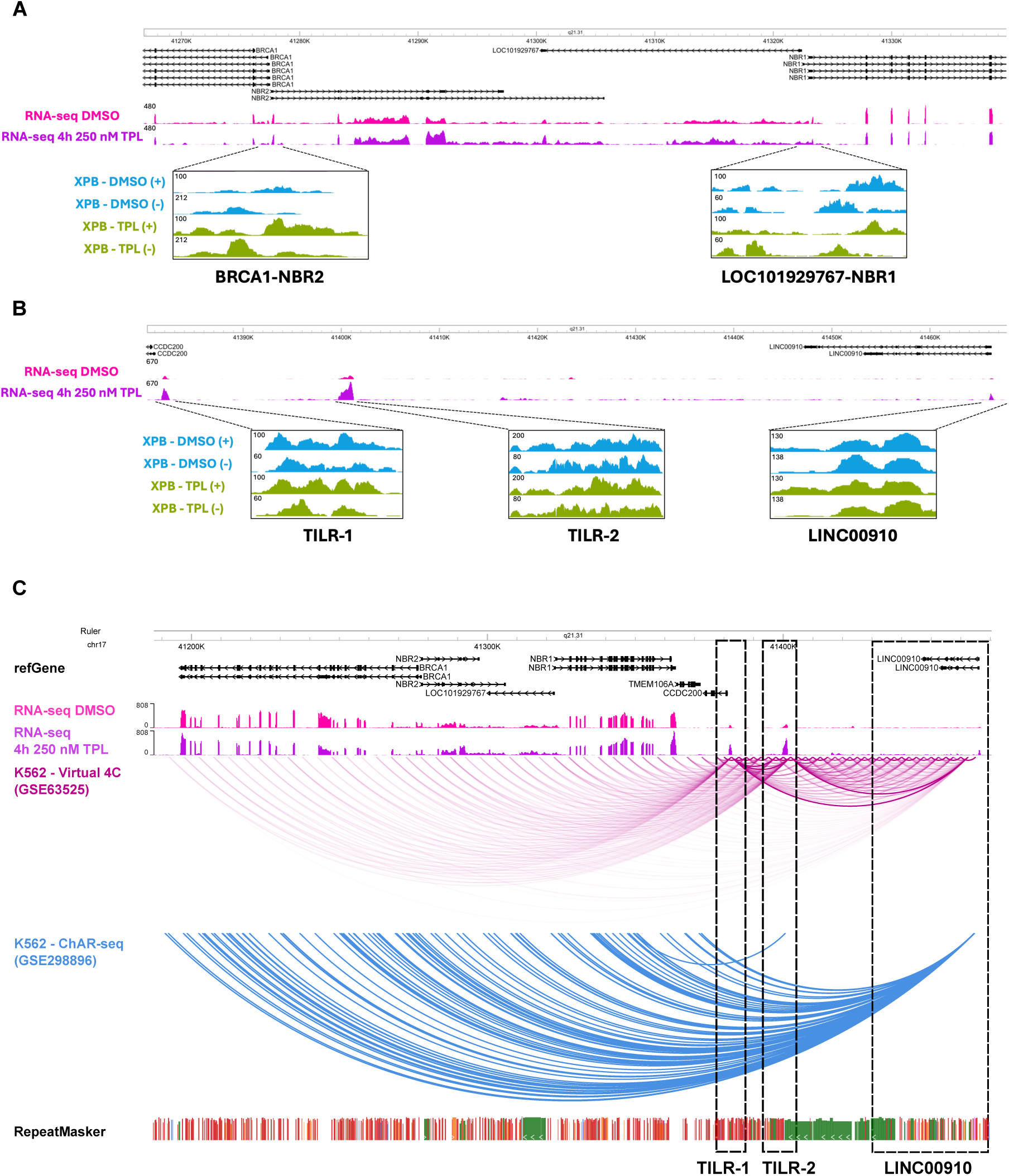
XPB occupancy on the *BRCA1* locus and in the *TILR-1*, *TILR-2* and *LINC00910* regions upon TPL and genomic interactions between the lncRNAs and the *BRCA1* locus. **A** Genome Browser image of the XPB occupancy measured by ChIP-exo in the regions corresponding to the *BRCA1 locus* in DMSO conditions or after 1 hour of 1 μM TPL treatment. **B** Genome Browser image of the XPB occupancy measured by ChIP-exo in the regions corresponding to *TILR-1*, *TILR-2*, and *LINC00910* in DMSO conditions or after 1 hour of 1 μM TPL treatment. The symbol (+) indicates the XPB occupancy at the sense strand, while the symbol (−) indicates the XPB occupancy at the antisense strand. (**C**) Genome Browser image of the chromosome 17 spanning the *BRCA1* locus and the adjacent locus where the two novel lncRNAs *TILR-1*, *TILR-2*, and *LINC00910* reside. Virtual 4C was performed using public Hi-C data from the GEO number accession GSE63525. ChAR-seq data was downloaded from the GEO number accession GSE298896. RepeatMasker data is shown below indicating the different classes of repeats (red = SINE, green = LTR, orange = LINE).

